# Coalescence and Translation: A Language Model for Population Genetics

**DOI:** 10.1101/2025.06.24.661337

**Authors:** Kevin Korfmann, Nathaniel S. Pope, Melinda Meleghy, Aurélien Tellier, Andrew D. Kern

## Abstract

Probabilistic models such as the sequentially Markovian coalescent (SMC) have long provided a powerful framework for population genetic inference, enabling reconstruction of demographic history and ancestral relationships from genomic data. However, these methods are inherently specialized, relying on predefined assumptions and/or limited scalability. Recent advances in simulation and deep learning provide an alternative approach: learning directly to generalize from synthetic genetic data to infer specific hidden evolutionary processes. Here we reframe the inference of coalescence times as a problem of translation between two biological languages: the sparse, observable patterns of mutation along the genome and the unobservable ancestral recombination graph (ARG) that gave rise to them. Inspired by large language models, we develop cxt, a decoder-only transformer that autoregressively predicts coalescent events conditioned on local mutational context. We show that cxt performs on par with state-of-the-art MCMC-based likelihood models across a broad range of demographic scenarios, including both in-distribution and out-of-distribution settings. Trained on simulations spanning the stdpopsim catalog, the model generalizes robustly and enables efficient inference at scale, producing over a million coalescence predictions in minutes. In addition cxt produces a well calibrated approximate posterior distribution of its predictions, enabling principled uncertainty quantification. We apply cxt to population genomic data from both humans and mosquitoes, highlighting the model’s ability to deal with the complexities of empirical data.

**Significance statement:** `cxt` is a language model for population genetics which introduces next-coalescence prediction as translation from observed mutations to coalescence times by modeling the coalescent with recombination as a conditional stochastic process. It learns implicit priors from `stdpopsim` and generalizes across both known and novel demographies. `cxt` generates millions of TMRCA estimates in minutes and samples well-calibrated posteriors for uncertainty quantification. A simple post-hoc correction aligns predicted diversity with the species mutation rate, ensuring robustness to novel evolutionary scenarios.

## Introduction

The stochastic process of genetic lineages tracing back to common ancestors over time gives rise to a complex structure known as the Ancestral Recombination Graph (ARG), with branches that merge through coalescence or divide through recombination (Kingman 1982; Hudson 1983; R. C. Griffiths and Marjoram 1997; R. C. Griffiths 1991). In the absence of mutations, these branches remain entirely concealed. However, mutations along the branches of the genealogy offer only a noisy, indirect window into this hidden ARG, carrying partial information about topology and the timing of events. Currently the field of population genetics is undergoing a revolution in our ability to infer the ARG that underlies a sample, and over the past few years we have obtained our first glimpses into reconstructions of whole genome genealogies (Kelleher et al. 2019; Speidel et al. 2019; B. C. Zhang et al. 2023; Deng, Nielsen, and Song 2024).

Inferring the ARG (or approximations of it) has long been a central goal in population genetics. One of the earliest attempts was by Griffiths and Marjoram 1996, who used Markov chain simulations of the coalescent process for ancestral inference. Subsequent work conceptualizing recombination as a point process along the genome (Wiuf and Hein 1999), shifted focus toward emphasizing the spatial aspects of recombination, and laid the groundwork for more efficient approximations of the coalescent with recombination such as the sequentially Markovian coalescent (SMC) (McVean and Cardin 2005). The SMC approximation has been pivotal in the development of numerous inference methods, beginning with approaches requiring only a single diploid genome to investigate demographic changes, such as PSMC (H. Li and Durbin 2011) (which effectively infers an ARG for two haploid samples). Since then, SMC-type methods have been extended in numerous ways including: to multiple genomes (Schiffels and Durbin 2014; Malaspinas et al. 2016), integration of site-frequency spectrum information (Terhorst, Kamm, and Song 2017), the inclusion of species-specific life-history traits (e.g. T. P. P. Sellinger et al. 2020; Korfmann et al. 2024; T. Sellinger, Johannes, and Tellier 2024), as well as algorithms that more efficiently scale as a function of sample size (Palamara et al. 2018; Schweiger and Durbin 2023; Terhorst 2024).

While the SMC approximation enables tractable likelihood-based inference for a restricted class of models, it becomes increasingly difficult to extend when key evolutionary processes, such as population structure, selection, or complex demographic events are introduced. Simulation-based inference (SBI) offers a flexible alternative by bypassing explicit likelihoods and instead learning mappings from data to model parameters directly from simulated datasets. Approximate Bayesian computation (ABC) is a widely used SBI framework in population genetics (e.g. Beaumont, W. Zhang, and Balding 2002), and has been successfully applied to a range of inference problems, including demographic history reconstruction (e.g. Boitard et al. 2016; Jay, Boitard, and Austerlitz 2019).

However, classical ABC methods rely on reducing data to low-dimensional summary statistics in order to mitigate the curse of dimensionality, often discarding information relevant to inference in the process. This limitation is particularly acute for sequence-based problems involving recombination, where genealogical information is distributed across long, high-dimensional haplotypic sequences. In such settings—exemplified by PSMC-like models—information loss during data compression can substantially limit predictive accuracy and robustness, motivating the development of inference approaches that operate directly on high-dimensional sequence data (see, however, (Min et al. 2025)).

A powerful alternative to standard ABC is to utilize deep learning (DL) for simulation-based inference, which has proven broadly effective across diverse domains (Schmidhuber 2022). In population genetics, DL-based methods have been successfully applied to a wide range of model-based inference tasks (as reviewed in Korfmann, Gaggiotti, and Fumagalli 2023), leveraging standard neural network architectures including fully connected networks (Sheehan and Song 2016; C. Battey, Ralph, and Kern 2020), convolutional neural networks (Kern and Schrider 2018; Flagel, Brandvain, and Schrider 2019; Torada et al. 2019; Sanchez et al. 2021; Whitehouse and Schrider 2023; Smith et al. 2023; Smith and Kern 2023; Saada et al. 2023), recurrent neural networks (Adrion, Galloway, and Kern 2020), and variational autoencoders (C. Battey, Coffing, and Kern 2021). However, most existing DL approaches in population genetics are trained for narrow, task-specific objectives, such as estimating individual demographic parameters, and often fail to generalize beyond the distribution of their training simulations. While domain adaptation strategies can partially mitigate this limitation (e.g. Mo and Siepel 2023), they do not fundamentally address the challenge of learning transferable representations of the underlying evolutionary process.

More recently, transformer-based language models (LMs) have been introduced that enhance prediction capabilities through the learning of stochastic processes, rather than scenario-constrained parameter estimation (e.g. Saada et al. 2023). Transformer LMs are perhaps best known in the context of pretrained generative models for human language such as GPT (Radford and Narasimhan 2018). These models are pretrained in that they are first trained on a “pretext” task, such as next token prediction, that requires the LM to learn context among observations (tokens) in a sequence locally and their relation to higher-level associations in the space of language. Modern large LMs are in part wildly successful due to their massive numbers of parameters (e.g. hundreds of billions) that enable them to effectively learn rich representations and flexible conditional prediction across contexts and thus to generalize. Here we describe a pretrained generative model for population genetics that leverages a small language model (10-20M parameters) to effectively translate from mutational patterns across chromosomes to coalescent time estimates. This opens the door to a generalizable (non-task specific) model paradigm: pretraining a flexible transformer on a diverse range of coalescent simulations, followed by fine-tuning for specific evolutionary tasks, if necessary.

In this work, we introduce `cxt`, the first language model for coalescent estimation to our knowledge. `cxt` is a decoder-only transformer model inspired by GPT-2 and designed for pairwise time to the most recent common ancestor (TMRCA) prediction —a core component of genealogical inference. In contrast to DNA language models such as GPN-MSA or Evo 2 (Benegas et al. 2025; Brixi et al. 2025) that are pretrained on the task of masked base prediction from a collection of sequenced genomes, our pretext task leverages synthetic data simulated under explicit population genetic models to predict the *next* pairwise TMRCA along a sequence. Rather than modeling DNA sequences itself, `cxt` aims to recover latent genealogical structure, providing a flexible and generalizable model for downstream inference. To enable this inference, we define a novel task, analogous to *next-token prediction* in language modeling, which we term *next-coalescence prediction*. In this formulation, the model predicts the next coalescence time along a sequence of previously predicted events, across discrete sequence space, conditioned on local mutation densities within a fixed context window. While this does not reconstruct full genealogical topologies, it enables a structured prediction task that translates between observed mutation patterns and pairwise coalescence times —akin to approaches such as PSMC or `SMC++`, the latter being the method to which `cxt` is conceptually a close neural analog, in the sense that prediction happens on a pivot (or distinguished pair), while sharing information across haplotypes through the SFS. Thus, the model offers a way to navigate between the observed distribution of mutation densities and the latent pairwise genealogical structure by means of a sequence of coalescence event prediction.

To evaluate this approach, we focused on three criteria: (1) **accuracy**—performance comparable to or better than recent theory-driven methods such as Singer and SMC++; (2) **robustness**—particularly under model misspecification and out-of-distribution scenarios; and (3) **capacity**—the ability to generalize across a diverse range of simulated evolutionary scenarios.

Using extensive simulations and applications to human and mosquito genomic variation data, we show that `cxt` provides competitive, state-of-the-art performance for inferring local coalescence times across recombining chromosomes, with efficient parallel GPU inference. The generative nature of `cxt` enables rapid sampling of an approximate posterior over TMRCA trajectories, providing uncertainty estimates that are well calibrated. Across a wide range of models, including nearly the full `stdpopsim` catalog, `cxt` maintains strong out-of-sample performance and can be further improved via targeted fine-tuning. Finally, we apply `cxt` to large-scale human and *Anopheles* population genomic data (Consortium et al. 2010; Auton and Salcedo 2015; 1000 Genomes Consortium et al. 2017; Clarkson et al. 2020), recovering known coalescent-time outliers in humans and time-resolved signals consistent with insecticide resistance dynamics in mosquitoes.

## Results

We begin by presenting a conceptual schematic that adapts the general language modeling paradigm to population genetics, and draw parallels with SMC-based coalescent modeling to define our next-coalescence prediction task (Figure 1). We then benchmark `cxt`’s accuracy against current fast inference methods, specifically `Singer` and `SMC++` (Deng, Nielsen, and Song 2024; Terhorst, Kamm, and Song 2017). Next, we evaluate the model’s cross-scenario generalization capabilities, training and evaluating using the nearly complete `stdpopsim` v0.2 and v0.3 catalogs across diverse species, demographic scenarios, and genetic maps (excluding only bacteria and two species with extreme recombination rates that preclude large-scale simulation and three models with ancient samples). We assess the global accuracy of TMRCA prediction by comparing true and estimated coalescence rates across simulations. Finally, we apply `cxt` to real genomic data, estimating coalescent times across two chromosomes from the 1000 Genomes Project, using pairs of haploid sequences of British ancestry as our focal samples and chromosome 2L of *Anopheles* from samples taken from Cameroon, Mali, Burkina Faso, Ghana and Uganda.

**Figure 1:**
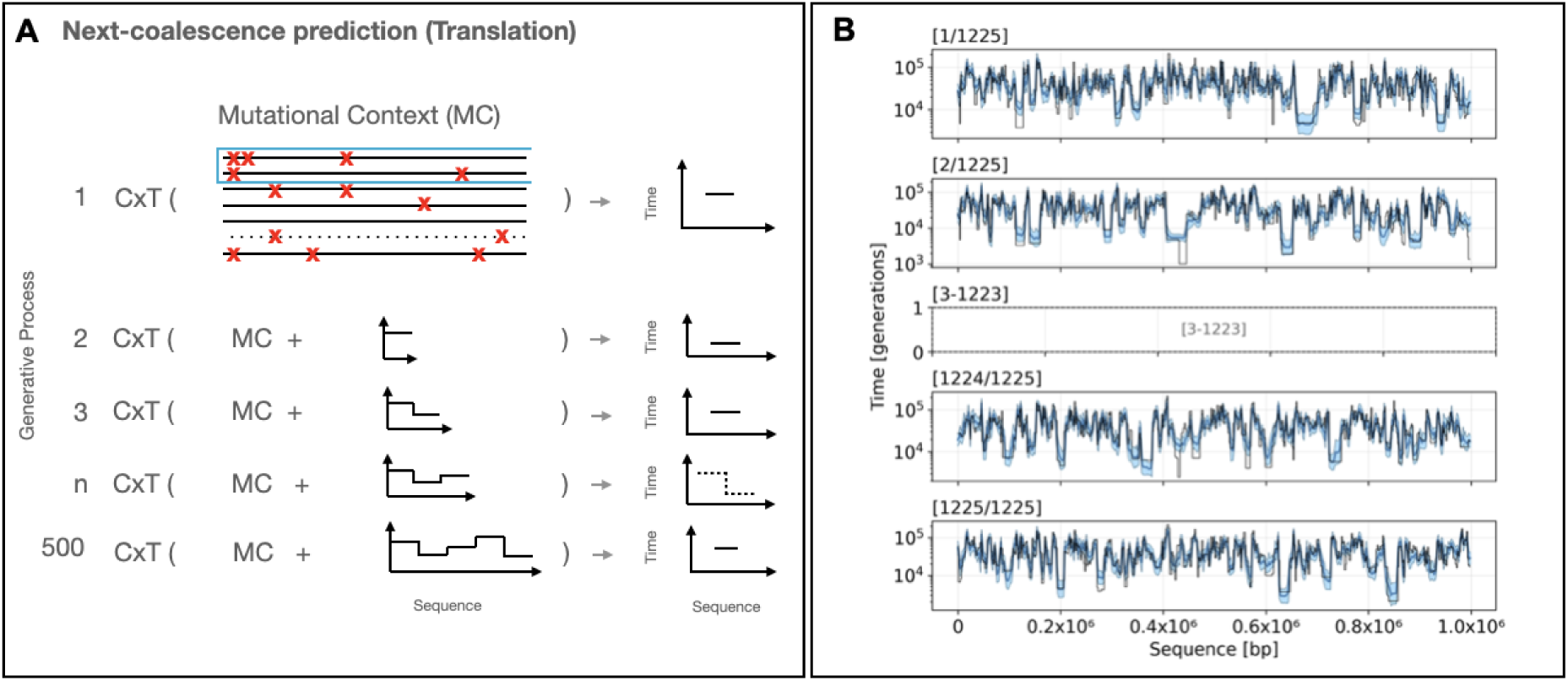
`cxt` introduces the notation of *next-coalescence prediction* (A). `cxt` is a language model that conditions on a chosen “pivot” haplotype pair and predicts the pair’s time to the most recent common ancestor (TMRCA) for each window. `cxt` ingests mutational tensors constructed using the pivot pair and site–frequency–spectrum (SFS) values computed in windows across a focal region. The model works autoregressively: after each window is predicted, that estimate is appended to the context and supplied to the next step, yielding a step-wise reconstruction of the entire pairwise coalescent history. Because every haplotype pair can be processed in parallel, the approach scales efficiently with available GPU memory. In panel (B) all 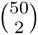 pairwise coalescence curves for a sample of 50 haplotypes were inferred simultaneously in under five minutes on a single NVIDIA A100 GPU. True TMRCAs are plotted in black, individual replicate predictions in light blue, and the prediction mean in dark blue.

### Fast and Accurate Inference

We developed `cxt`, a decoder-only transformer architecture adapted for the task of inferring local coalescence times from mutation patterns across recombining genomes. Unlike language models that embed discrete tokens, `cxt` projects continuous mutation densities into a latent space using feed-forward networks, followed by attention-based contextualization. Rotary positional embeddings encode physical distance along the genome, allowing the model to capture locality in genealogical relationships. Trained using a next-coalescence prediction objective, `cxt` captures the stochasticity of evolutionary processes while achieving rapid and accurate inference.

We show a descriptive schematic of the model in Figure 1A. `cxt` follows a general language modeling paradigm in which the model predicts a discretized coalescent time for each genomic window based on preceding predictions and the local mutational context. Specifically, we divide the genome into fixed-size positional windows and discretize coalescent times into bins on a logarithmic scale. Given a sequence of ground truth discretized pairwise TMRCA values `E` = (`e`_1_`, e`_2_`, . . ., e`*_n_*), the model autoregressively learns the conditional distribution:

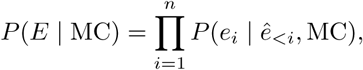

where 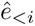 represents the model’s stochastic predictions for the preceding windows and MC denotes the mutational context (the observed data) across all windows. Concretely, the mutational context in a given window is the site frequency spectrum calculated across all samples, partitioned into the contributions from sites that are heterozygous and sites that are homozygous in the pivot pair.

At each genomic window i, the model outputs a probability distribution over discretized TMRCA bins:

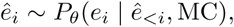

with `P`*_θ_* parameterized by a neural network with weights *θ*. The training objective is to maximize the log probability of the true sequence given the (preceding) predicted sequence:

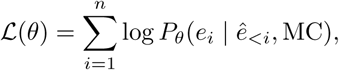

where *e_i_* denotes the ground truth label in window *i*, and 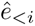 are the model’s own predictions up to (but not including) window *i*. Predictions are then centered around an intercept (log expected total diversity), which is sampled from the posterior distribution conditional on observed total diversity assuming a Poisson model of mutation. This objective encourages the model to learn a mapping from local mutation patterns to genealogical structure, enabling accurate sequential inference of coalescent events across the genome.

Figure 1B shows a concrete example of the model’s output. Leveraging our GPU-enabled method, we infer pairwise coalescence times for all 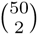 pairs of a sample of 50 haploid chromosomes in parallel by placing them in a batch of size 1225 (although note this batch size is dependent on GPU memory, and can be adjusted to available GPU memory). We then run the generative process multiple times, typically 15 replicates, and average the results to obtain stable and robust estimates. This generative process provides samples from an approximate posterior distribution of pair coalescence times across the sequence (where the prior is implicit via the simulated training set; see Calibration of approximate posteriors). In Figure 1B, we apply our `cxt` to a constant population size scenario with a population size of 2 × 10^4^ and approximately equal mutation and recombination rates (see Datasets and Training Description below). In that figure true coalescent times are shown in black, the replicate predictions in light blue, and the average prediction in dark blue. In this simple setting, where the model is well-specified, `cxt` produces highly accurate predictions. Importantly, `cxt` can predict coalescent times for *all* pivot pairs concurrently in approximately five minutes of computation on an NVIDIA A100 GPU with 80GB memory.

### Benchmark Comparisons

To evaluate cxt’s performance, we benchmarked its accuracy against two approaches Singer and SMC++ (Deng, Nielsen, and Song 2024; Terhorst, Kamm, and Song 2017), selected for their accuracy and broad adoption, respectively.

We first consider the performance of our “narrow model”—a `cxt` model trained on a constant-sized population (*N_e_* = 2 × 10^4^) with equal mutation and recombination rates (see Datasets and Training Description). In Figure 2, we aim to demonstrate two main results. First, we show the performance of the narrow model (**cxt**-narrow, top-left) on its original training domain—constant population size with roughly equal mutation and recombination rates—which yields a mean squared error (MSE) of 0.2447. As expected, **cxt**-narrow’s predictive accuracy declines when evaluated outside this domain. The top-middle panel shows results under a fluctuating population-size model (the sawtooth scenario; MSE 0.6431); thus, model misspecification degrades performance. We can potentially mitigate this behavior by training `cxt` on a wider range of parameters and demographic histories. To do so, we introduce a “broad model”—**cxt**-broad (top-right)—trained on a substantially wider range of scenarios (see Datasets and Training Description). **cxt**-broad, because it can accommodate demographic variation, achieves improved accuracy on the sawtooth model (MSE 0.1807).

**Figure 2:**
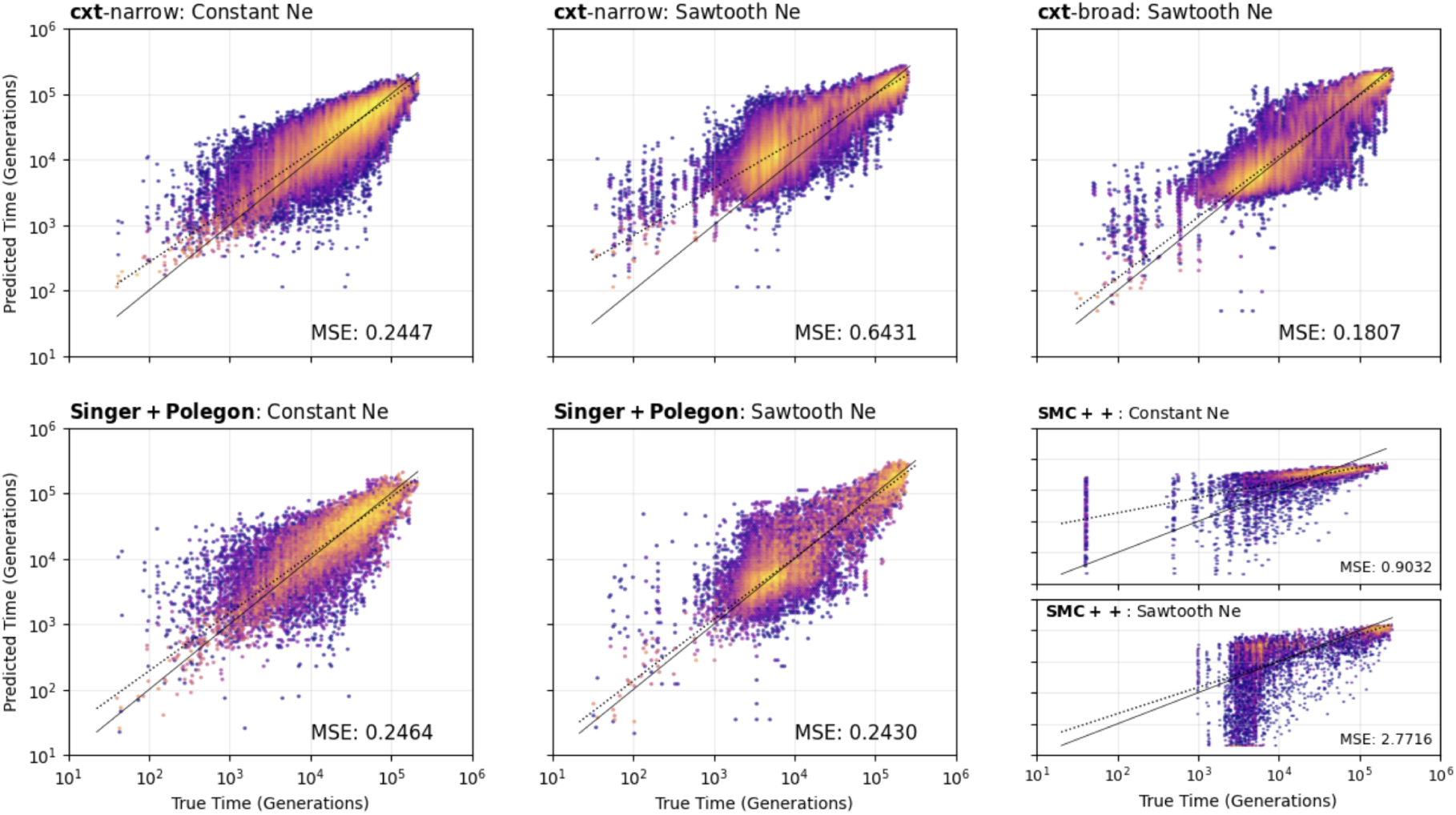
True versus predicted coalescence times for three inference approaches across two demographic scenarios: a constant population size and a fluctuating “sawtooth” demography. **Top row**: `cxt-narrow` evaluated on the constant-size scenario (left) and on the sawtooth scenario (middle), followed by `cxt-broad` evaluated on the sawtooth scenario (right). The broad model—trained on a wider range of demographic histories—achieves substantially improved accuracy. **Bottom row**: `Singer+Polegon` evaluated on the constant-size (left) and sawtooth (middle) scenarios. The two rightmost panels show `SMC++` applied to the constant-size and sawtooth scenarios, respectively. Although `SMC++` is primarily designed for population-size inference, we include its decoding-based coalescence-time estimates for completeness. Mean squared errors (MSE) for each panel are reported within the plots.

Singer, combined with its dating-refinement algorithm Polegon (**Singer+Polegon**), attains comparable accuracy on the constant-size scenario (Figure 2 bottom-left; MSE 0.2464) and slightly reduced accuracy under the sawtooth model (MSE 0.2430). **SMC++**, which is primarily designed for demographic history inference with TMRCA decoding provided as part of its composite-likelihood framework, yields higher MSE values for these simulations (0.9032 and 2.7716), reflecting its emphasis on population-size reconstruction rather than fine-scale local genealogy prediction.

As `cxt` is trained at a specific sample size, the user has a few options should they want to infer coalescent times in a new sample of different size: 1) they could retrain the model starting from a new set of simulations that condition on their sample size, 2) if the sample size is larger than the trained model, they could subsample down to the appropriate size, or 3) if the sample is smaller than the trained model we have created a light-weight adapter, which quickly fine-tunes a larger sample size trained `cxt` model to a smaller sample size. In Supplementary Figure S4, we show representative results from the adapter applied to samples of size *N* = 5 diploids from a `cxt` model trained at sample size *N* = 25 diploids, and then used for inference of a constant sized simulation (left panel) or the sawtooth simulation (right panel).

Our results demonstrate that the language modeling approach implemented in `cxt` is highly competitive with the most recent theory-driven methods, whether ARG-based (as in `Singer`) or SMC-based (as in `SMC++`). A full summary of error estimates is provided in Table 1 and Figure 2.

**Table 1:**
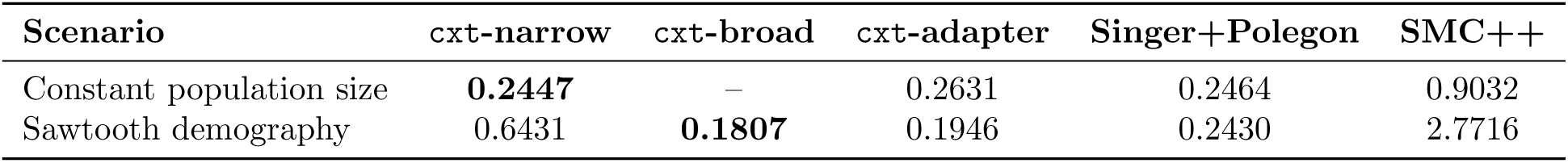
Mean squared error (MSE; lower is better) for three approaches across two demographic scenarios. We report results separately for the narrow and broad variants of `cxt` (that `cxt`-adapter uses the broad model as its basis) alongside **Singer+Polegon** and `SMC++` reflecting training on a single demographic regime (constant size) versus a fluctuating demographic history (sawtooth), respectively.

| Scenario | cxt-narrow | cxt-broad | cxt-adapter | Singer+Polegon | SMC++ |
| --- | --- | --- | --- | --- | --- |
| Constant population size | **0.2447** | – | 0.2631 | 0.2464 | 0.9032 |
| Sawtooth demography | 0.6431 | **0.1807** | 0.1946 | 0.2430 | 2.7716 |

### Computational Efficiency and Scaling

We compared wall-clock runtime for `cxt`, SMC++, and `Singer` across increasing numbers of inferred pairwise coalescence trajectories (Supplementary Figure S6). Because these methods differ fundamentally in how inference is performed, runtime is evaluated separately from statistical accuracy. `SMC++` achieves fast local decoding, but total runtime is dominated by preprocessing and composite-likelihood optimization, both of which increase with dataset size. Note that `SMC++` also assumes unphased samples—a constraint that could be addressed by restructuring the VCF file itself, but which we omit here, as runtimes are already large. In contrast, `cxt` performs fully amortized inference: after training, runtime scales approximately linearly with the number of inferred pairs and shows near-linear speedups with additional GPUs. Runtime is largely insensitive to recombination rate, enabling predictable performance across parameter regimes, but preprocessing overhead is noticeable when going beyond 50 samples. `Singer` shows competitive accuracy but substantially higher runtime, with sensitivity to recombination parameters reflecting its reliance on explicit genealogical sampling. Overall, these results highlight a qualitative difference in computational scaling: while likelihood-based and MCMC-based methods incur increasing cost as genealogical complexity grows, `cxt` shifts this cost to training, enabling fast inference at application time, while as of now being constrained by the pairwise binomial scaling with the number of samples.

### Towards a generalizable deep learning model

In the previous section, we focused primarily on the setting in which `cxt` is trained on a single demographic model (`cxt`-narrow), thereby learning directly from simulated data, while also providing a preview of `cxt`-broad’s potential. The central advantage of a language-modeling approach is its capacity—through many learnable parameters— to move beyond narrowly specified generative assumptions by conditioning its behavior on a rich set of mutational and genealogical contexts. In this sense, `cxt` functions as a flexible conditional density estimator: for a given context, it produces a corresponding predictive distribution. We assess this directly by comparing inferred marginal coalescence-time distributions across species.

In Figure 3, we report inferred coalescence time distributions across nearly all `stdpopsim` v0.2 species. For each species, we simulate under its published demographic model when available, and otherwise assume a constant population size. Results from simulations using fine-scale recombination maps are given separately in Supplementary Figure S7. All results shown here are based on the `cxt`-broad model (see Datasets and Training Description), which was trained across the full set of scenarios. Importantly, the evaluations in Figures 3 and S7 are performed on newly generated simulations that were not included in training, eliminating any opportunity for data leakage. Across nearly all species, the inferred distributions (blue) closely match the true distributions (black).

**Figure 3:**
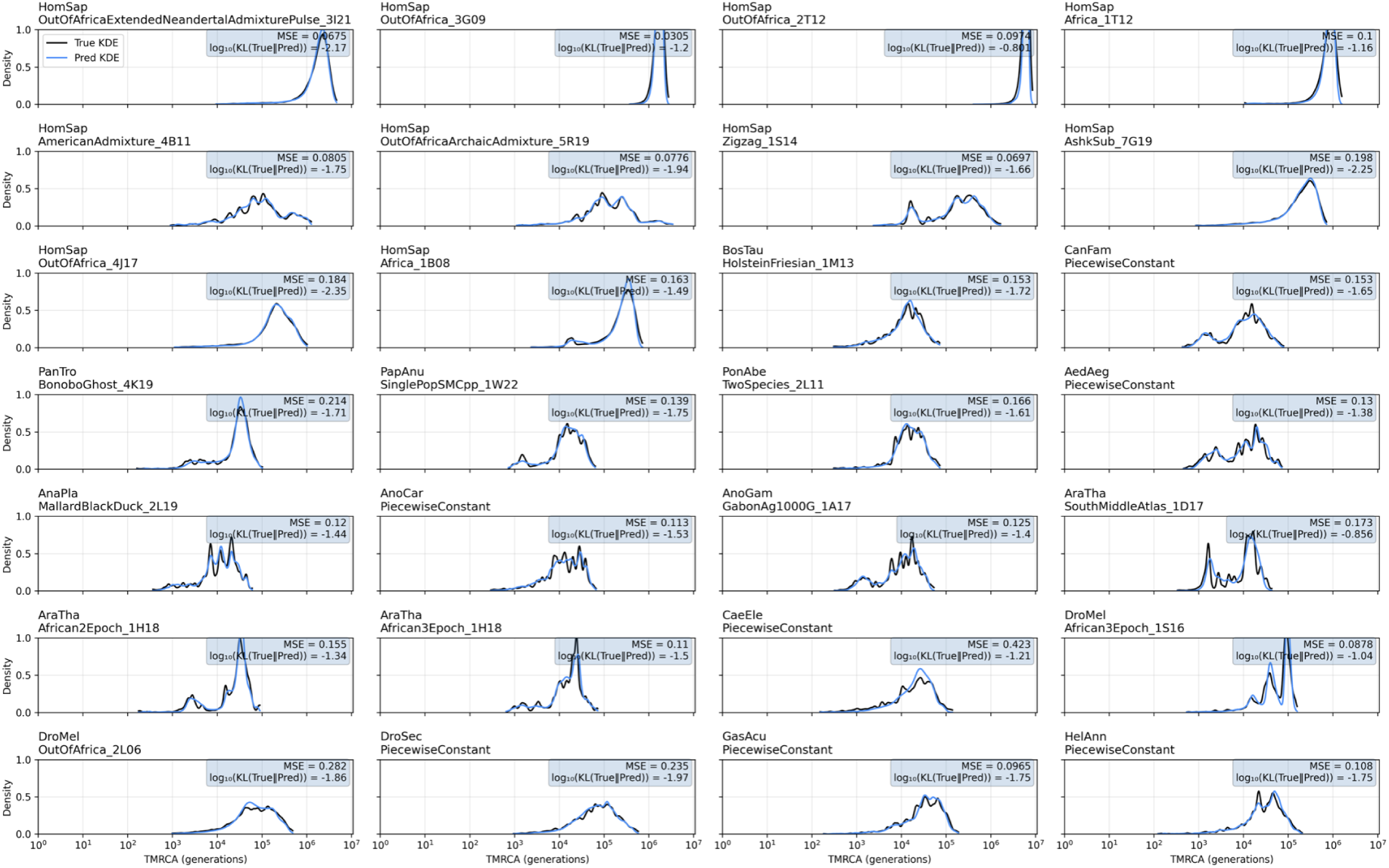
Evaluation of marginal coalescence distribution inference (blue line) against the true distribution (black line), probing the model’s capacity to distinguish among many scenarios based on context alone. The broad model’s ability to infer a `stdpopsim` v0.2 coalescence distribution is tested, which includes diverse mutation and recombination rates, as well as demographic scenarios. This distribution is obtained by aggregating pairwise inferences and visualizing the result as a kernel density for each scenario. All results shown are from new simulations, not included in the original training dataset. No genetic map was used for the inference shown here; the model itself was trained with and without genetic maps. For inferences with an underlying genetic map see the corresponding Figure S7.

To promote generalization across demographic and rate regimes, we made a deliberate design choice during training: we keep the window size fixed at 2 kb while varying *N_e_* over several orders of magnitude. In principle, one might expect the window size to scale with the effective population size, since the expected numbers of mutations and recombination events per window scale with *N_e_*. However, by intentionally not adapting the window size, the model encounters windows that can be mutation-dense or mutation-sparse depending on the underlying *N_e_*. This encourages the model to learn *relative* changes in local coalescent structure rather than relying on absolute mutation counts; any global offset can be corrected during inference by calibrating to the mutation rate.

A related consideration is how well the mean coalescent time within a window can be estimated. As the window size increases, estimation becomes easier: mutation counts rise, diversity increases, and sampling noise decreases. However, larger windows also average over spatial variation in genealogical structure, attenuating signals of population-size change, population structure, and other localized effects.

We illustrate this trade-off in Figure S5. Using 2 kb windows (left; MSE 0.1096) predicted coalescence times closely track the truth for the `stdpopsim` *AnoGam* model. In contrast, using much smaller 0.2 kb windows (middle; MSE 0.5727) substantially increases difficulty: more frequent recombination induces greater heterogeneity among local genealogies, while fewer mutations per window increase variance in the observed signal. Despite the broad model’s reasonable performance in this setting, we further fine-tune it on `stdpopsim` species with *N_e_* *>* 10^5^ (Figure S5 right). This yields the `broad (w200)` model and further reduces prediction error.

### Out-of-sample tests for the broad model

During this project, `stdpopsim` v0.3 was released, expanding the catalog to include additional species (*S. scrofa, R. norvegicus, P. sinus, M. musculus*, and *G. gorilla*). We leveraged this release to assess the generalization ability of our broad model, testing performance *without* retraining. Because these species were not present in `stdpopsim` v0.2, they constitute genuinely out-of-sample test cases for the broad model.

The v0.3 release includes species spanning a wide range of effective population sizes, from very large populations, such as *S. scrofa* (*N_e_* = 270000) and *R. norvegicus* (*N_e_* = 124000), to much smaller populations, such as *P. sinus* (*N_e_* = 3500). Most of these new species also exhibit recombination rates that exceed mutation rates. For example, in *P. sinus* we have *m* = 5.83 × 10*^−^*^9^ and *r* = 1 × 10*^−^*^8^ per bp per generation (*r/m* = 1.715). A notable exception is *G. gorilla* for which r = 1.193 × 10*^−^*^8^ and m = 1.235 × 10*^−^*^8^, yielding r/m = 0.966.

Results from this out-of-sample evaluation are shown in Figure 4. Notably, `cxt` generalizes well, even when applied to simulations generated under previously unseen regimes of recombination and mutation rates, population sizes and demographic structure. Across these v0.3 species, `cxt` performs on par with `Singer+Polegon` and substantially better than the `SMC++` decodings.

**Figure 4:**
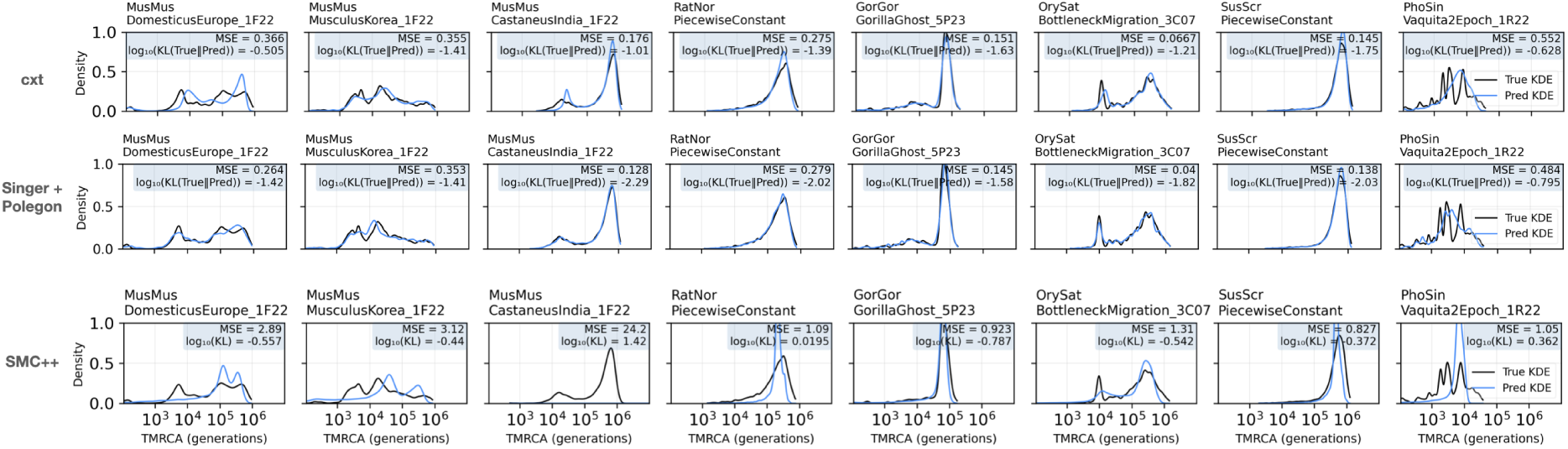
Out-of-sample evaluation of the broad model on `stdpopsim` v0.3. Each panel shows inferred marginal coalescence distributions (dashed) against true distributions (shaded). We simulated data from the novel species / demographic histories added in `stdpopsim` v0.3, aggregating pairwise inferences into kernel density plots. All results are from simulations previously unseen during training (Top-Row: `cxt`; Middle-Row: `Singer+Polegon`; Bottom-Row: `SMC++`).

### Generalization across parameter regimes

A central goal of this project was to build a model that spans the diversity of scenarios represented in`stdpopsim`, motivated by the idea that as `stdpopsim` expands, we can incrementally incorporate new species into training. However, when applying `cxt` to data from a new species—with potentially different mutation and recombination rates, effective population size or population structure— we expect inference accuracy to depend on how similar that species is to the regimes represented in the training data.

In Supplementary Figure S11, we assess the model’s ability to interpolate and extrapolate beyond the training distribution with respect to mutation and recombination rates, holding *N_e_* fixed at 20,000. We define a grid of mutation and recombination rates chosen to encompass most `stdpopsim` species (panel **A** in Figure S11); the distribution of species across this grid is shown in Figure S11 (**B**). Figure S11 (**A**) also highlights a range of recombination-to-mutation rate ratios Figure S11 (**B**), while Figures S11 (**C**) and (**D**) report performance over the grid in terms of MSE and KL divergence, respectively. As expected, error is highest in the low-mutation, high-recombination region (bottom right of Figure S11 (**C**)) where the signal-to-noise ratio is the least favorable for TMRCA inference. Figure S12 provides representative examples of TMRCA predictions from the four corners of the grid. Together, these results show that `cxt` can generalize beyond the specific regions of parameter space represented in the training data.

### Calibration of approximate posteriors

While the results above focus on point accuracy, `cxt` also samples TMRCA trajectories from an approximate posterior, enabling uncertainty quantification in addition to producing point estimates (e.g. the posterior mean). We therefore tested empirically whether the marginal approximate posteriors are (i) well-calibrated in the sense of providing correct frequentist coverage of the true TMRCA, and (ii) consistent, in the sense that the posterior concentrates as the amount of mutational information increases.

To assess these properties, we first simulated 1,000 completely independent pivot pairs under three demographic scenarios, including one drawn from the out-of-training `stdpopsim` v0.3 release (*Oryza sativa*). For each pair, we computed posterior intervals for the TMRCA in each 2 kb window using 100 `cxt`-sampled trajectories. For an exact posterior, an interval containing a fraction *α* of posterior mass contains the true value with probability *α*. We therefore evaluate calibration by computing, across windows, the empirical fraction of true TMRCAs that fall within posterior intervals of varying nominal mass. Second, we evaluate consistency by measuring how the average posterior variance changes as a function of mutation rate, scaled by 0.5, 1, and 2 relative to the species-specific mutation rate reported in the `stdpopsim` catalog. Both of these measures are calculated with respect to the position within the full 1Mb prediction window, since we expect performance to degrade near the boundaries where the local mutational context is truncated.

Across three distinct parameter regimes, we find that `cxt` produces generally well-calibrated posterior samples across most positions within the 1 Mb prediction window (Supplementary Figure S8, top row). These approximate posteriors are typically slightly over-concentrated high nominal coverage (e.g., intervals containing 95% of the posterior mass contain the true TMRCA roughly 92% of the time), but are otherwise remarkably stationary: the empirical coverage declines appreciably only in the final 2 kb window. This drop appears to be driven by a tendency of the trained model to predict occasional jumps in the last window, which can be addressed in practice by omitting that window prior to centering the predictions. Consistent with posterior concentration, increasing (decreasing) mutational density, causes the posterior intervals to contract (expand) reflecting increased (decreased) information (Supplementary Figure S8, bottom row). As expected, windows near the boundaries also exhibit greater uncertainty, consistent with truncation of the local mutational context.

Together, these findings indicate that the approximate posteriors produced by `cxt` provide reliable uncertainty quantification for predicted coalescence times, across demographic scenarios and mutational densities. This is a key strength of the method, particularly because posterior sampling is so computationally efficient.

### Demography estimation through coalescence rates

Estimating pairwise coalescence rates provides a direct window into changes in effective population size through time, perhaps most famously exemplified by the original `PSMC` model of Li and Durbin (2011). To evaluate our ability to recover effective population size trajectories from `cxt` predictions, we simulated data under demographic models for *H. sapiens*, *B. taurus*, and *A. thaliana*. From these simulations, we estimated pairwise coalescence-time distributions and converted them into coalescence-rate curves, which we then mapped to estimates of historical effective population size (Figure 5; see Methods).

**Figure 5:**
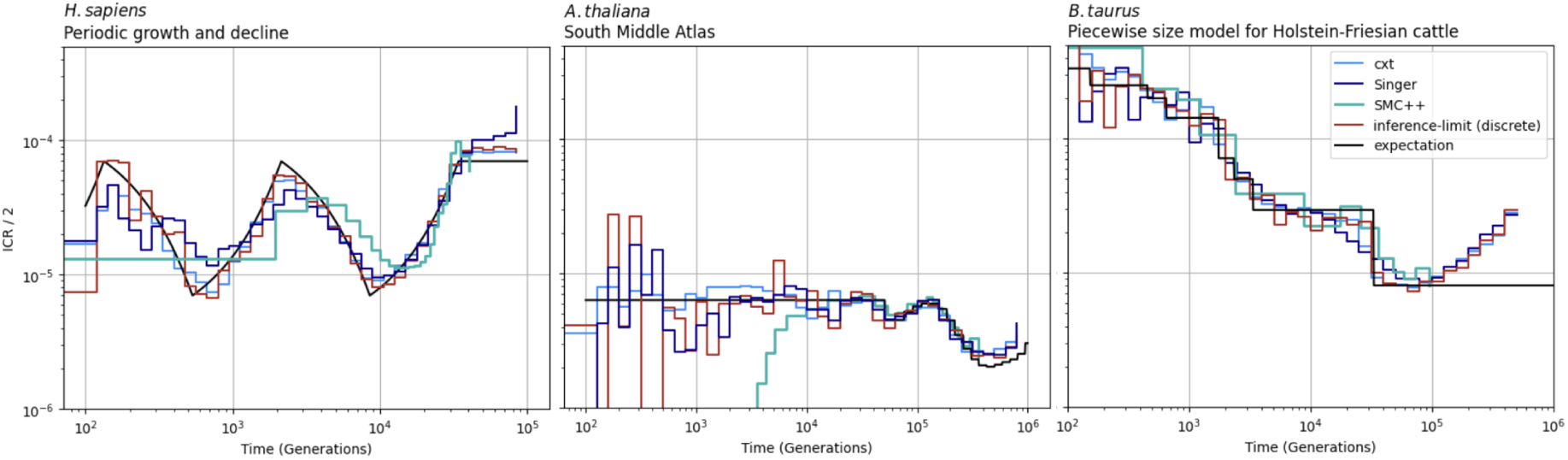
Inverse-instantaneous coalescence rate calculation for the piecewise-constant demography of (**A**) *H. sapiens*, (**B**) *B. taurus* and (**C**) *A. thaliana*. The inference of pairwise-coalescence events leads to the implicit inference of demography estimates through the marginal coalescence distribution assuming coalescence occurs as a Poisson process (see methods). For each scenario 10 Mb and 25 diploid samples have been used to achieve resolution throughout the specified time-windows.

Figure 5 shows five curves. The black *expectation* line reflects the true coalescence rate implied by the generative model. We also report an *inference limit* in red, constructed by sampling the true TMRCAs every 2 kb across the sequence and then estimating coalescence rates using the same procedure applied to `cxt`, `Singer`, and `SMC++`, thereby providing an upper bound under matched discretization. The remaining curves show the corresponding estimates from `cxt`, `Singer`, and `SMC++`.

Overall, effective population size trajectories inferred from cxt and Singer closely track the truth at a genome-wide scale capturing the major epochs of growth/bottlenecks, whereas SMC++ shows larger deviations at recent times; this gap may reflect limited information in the current setting and could plausibly be reduced by incorporating additional data.

We also inferred the cross-coalescence rate (i.e., the rate of coalescence between lineages drawn from different demes) under the two-population `OutOfAfrica 2T12` model from `stdpopsim`. Generally, cross-coalescence curves are informative about the timing of divergence and subsequent gene flow between populations within a species. Figure S10 summarizes these results. The left panel shows the cross-population rate estimated from one African and one European haplotype, whereas the middle and right panels show the within-population rates for African and European samples, respectively. The cross-population curve (left) closely follows the expected trajectory. In contrast, the within-population curves exhibit a modest deviation for `cxt`.

A likely explanation is the way structured models were represented during training: for each simulated instance, 50 samples were drawn with random population assignments. As a result, training examples rarely contained only a single population, which may bias within-population rate estimation. This could potentially be mitigated by modifying the training scheme for structured models, for example by explicitly including single-deme sampling configurations.

It is worth noting that the demographic inferences shown in Figures 5 and S10 are based on relatively short simulated sequences. For computational efficiency, we restrict these demonstrations to 10 Mb of simulated genome. Because shorter sequences increase the variance of recent-time coalescence-rate estimates by truncating the longest tracts of recent ancestry, we expect that applying the same procedure to full-length chromosomes would yield substantially improved accuracy. Together, these results show that `cxt`’s local TMRCA predictions can be aggregated into accurate genome-scale summaries of demography, with performance expected to improve further at realistic chromosome lengths.

### Application to Human 1000 Genomes and Ag1000G Mosquito data

Having established the accuracy and robustness of `cxt` across a broad range of simulated scenarios, we next apply it to empirical genomic data: human variation from the 1000 Genomes Project (Consortium et al. 2010) and mosquito genomes from the Ag1000G consortium (1000 Genomes Consortium et al. 2017).

We begin with the human dataset, revisiting the canonical selective sweep at *LCT* as a validation benchmark, and then turning to more recently dated regions of the *HLA* locus. In the latter case, our inferred TMRCA patterns are broadly consistent with previously reported old genealogical ages, though we infer slightly younger than `Singer+Polegon`.

We then analyze the insecticide-resistance loci *Rdl* in *Anopheles gambiae*. This locus has been highlighted using summary-statistic approaches, including Garud’s `H`_1_`/H`_12_, but a clear time-resolved interpretation of the underlying sweep dynamics has remained elusive. Using `cxt`, we estimate fine-scale coalescent times across five African populations—Burkina Faso, Mali, Cameroon, Ghana, and Uganda—while addressing three technical challenges characteristic of Ag1000G data, and we compare our mosquito estimates to `Singer+Polegon`:

1. **Large effective population sizes**, which yield deep genealogies and correspondingly large scaled mutation and recombination rates;
2. **Small sample sizes** in Ghana (< 25 diploid individuals); and
3. **Substantial missing-data variation** across genomic regions

To begin this initial validation in humans, we selected 50 individuals of British descent (GBR), and estimated coalescent times for 25 focal pairs, holding out the remaining n = 48 samples as the AFS input to cxt.

We choose 25 focal pairs because this sample size allows all chromosome-wide comparisons to fit in a single batch on an A100 GPU, enabling rapid whole-chromosome inference. The top panels of Figure 6 show chromosome-wide TMRCA landscapes for chromosomes 2 and 6. As an additional validation, we compared inferred TMRCAs against the number of recombination events predicted by `cxt` (Fig. S13), recovering the expected inverse relationship between genealogical age and segment length.

**Figure 6:**
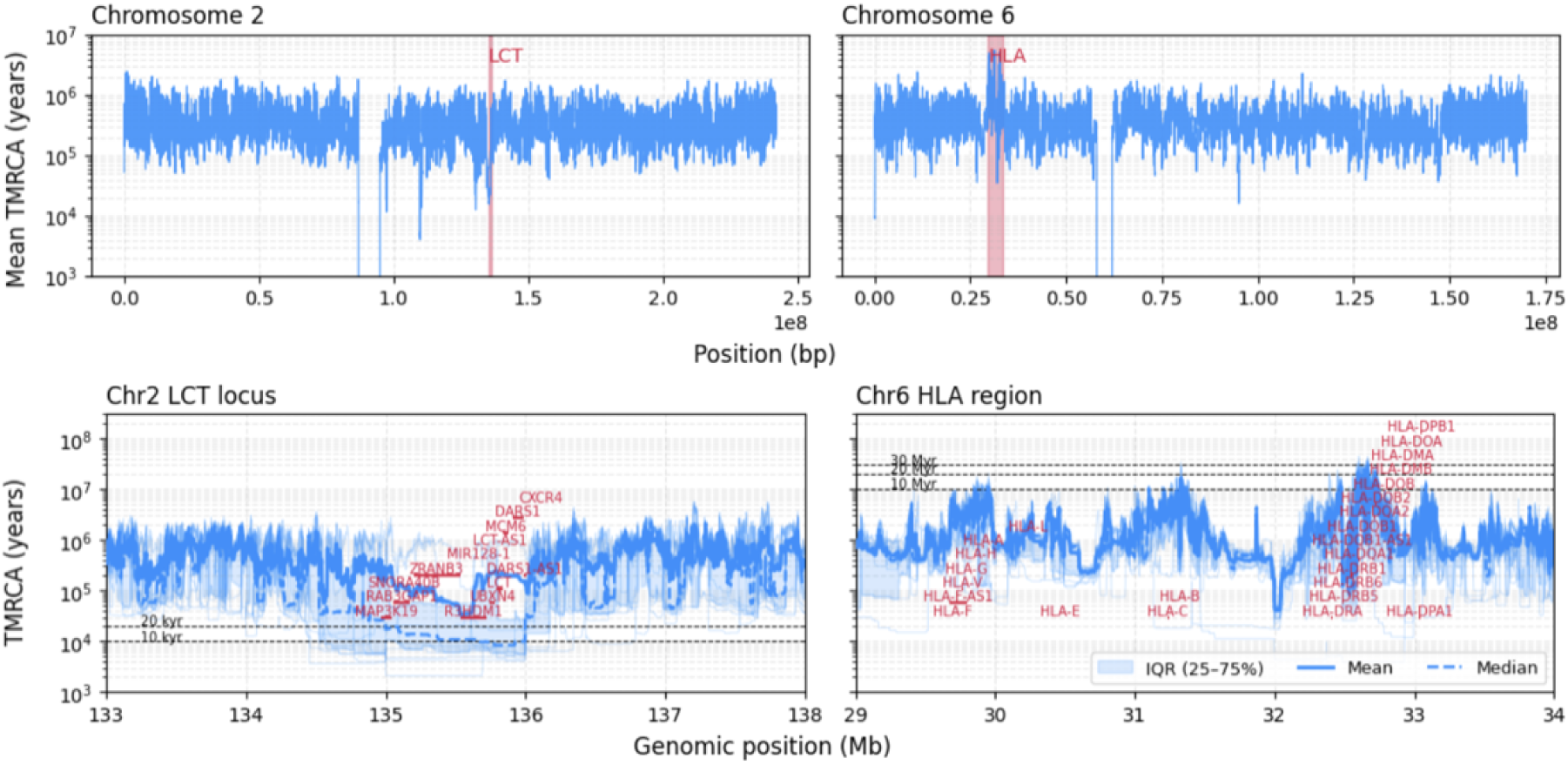
Inference of coalescent times from 25 focal pairs of British individuals of the 1000 Genomes project. The top-left panel shows chromosome 2, and the top-right panel shows chromosome 6. The LCT region (bottom-left) and HLA region (bottom-right) are zoomed respectively. Each pivot pair has been inferred 15 times to get a reliable mean, median and variance estimates. Additionally, light blue lines show the respective inferences of the 25 pairs.

The bottom panels in Figure 6 zoom in on two canonical regions: the *LCT* locus on chromosome 2 and the *HLA* region on chromosome 6. At *LCT*, we recover markedly reduced coalescent times as expected under the well-characterized recent selective sweep at this locus. `cxt` captures this local trough cleanly. Around *LCT*, estimated coalescent times slightly above 10 kya are consistent with the origin of the sweeping haplotype, and the variation across focal pairs highlights local heterogeneity.

The *HLA* region provides the complementary pattern: exceptionally deep genealogical structure shaped by long-term balancing selection. The extended peaks inferred by `cxt` (bottom-right) match long-standing expectations for this locus and support the conclusion that the model captures extreme TMRCA variation. Consistent with this, we replicate the multi-peaked ancient structure reported by (Deng, Song, and Nielsen 2025), with several genes reaching TMRCAs on the order of tens of millions of years, one of the strongest known signatures of long-term balancing selection.

To investigate selective dynamics at the *Rdl* locus across multiple *A. gambiae* populations, we constructed a comparison panel (Figure S9) comprising (i) coalescent-time landscapes inferred by `cxt` (left column), (ii) corresponding estimates from `Singer+Polegon` (middle column), and (iii) SMC++ decodings (right column). Analyzing Ag1000G data poses two practical challenges. First, large effective population sizes imply high *N_e_*-scaled mutation and recombination rates, which increase genealogical heterogeneity and the number of recombination events, substantially slowing MCMC-based approaches such as `Singer+Polegon`. Second, missingness varies widely across genomic regions, requiring methods that can accommodate irregular observation patterns (density of missing data is shown beneath the upper and lower panels as a track in Supplementary Figure S9). To improve readability in the main text, we restrict these figures to `cxt` (Figure 7); full comparisons with `Singer+Polegon` and other approaches are shown in the Supplementary Material, where these analyses are presented in detail.

**Figure 7:**
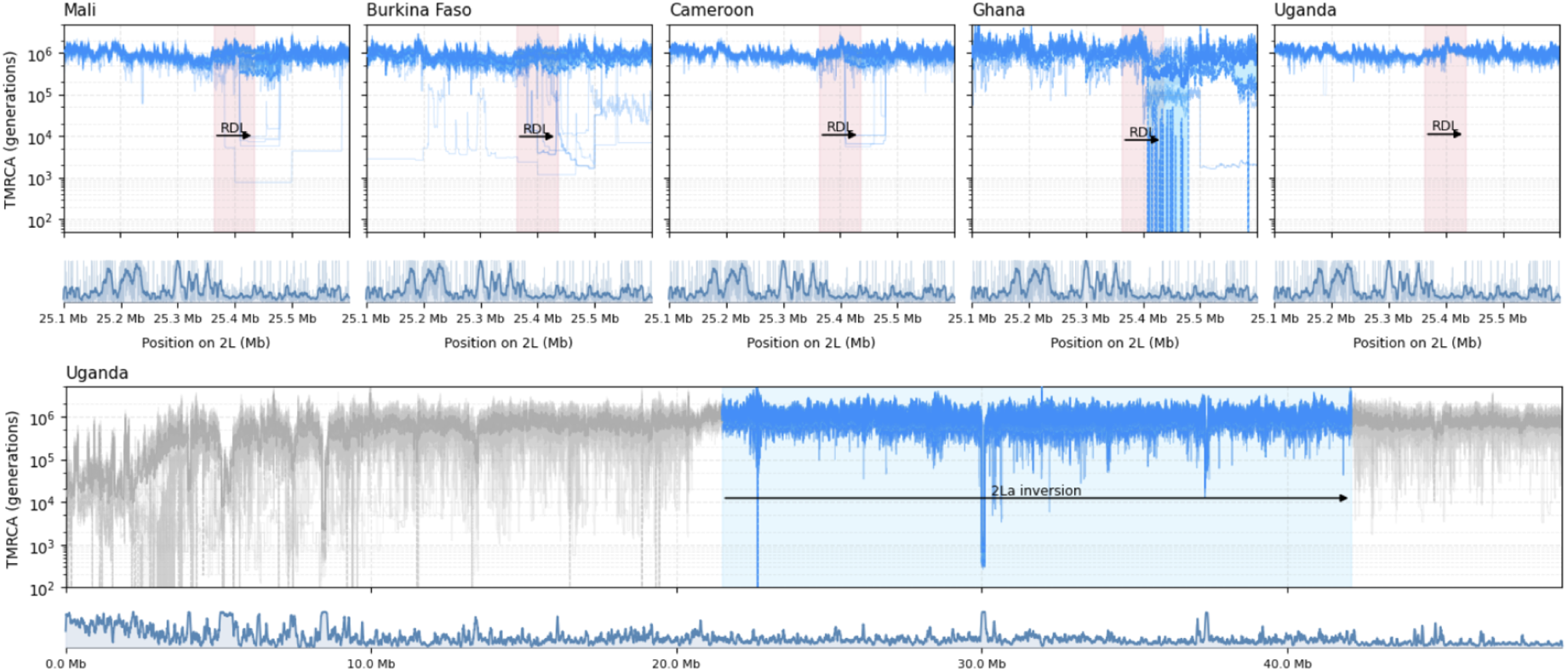
Inference of coalescent-time landscapes in the Ag1000G *A. gambiae* dataset across five African populations using `cxt`. In the upper panel, we analyze the *Rdl* region for Burkina Faso, Mali, Cameroon, Ghana, and Uganda. For these countries, we infer coalescent times for 25 focal haplotype pairs per population; for Ghana, we analyze five diploid individuals. Light-blue curves indicate per-pair coalescent-time trajectories inferred by `cxt`, with the mean trend shown in darker blue. The lower panel shows genome-wide coalescent-time landscapes across the entirety of chromosome 2L for Uganda samples. Missing data density is shown beneath both the regional and genome-wide panels as a genome browser track.

**Figure 8:**
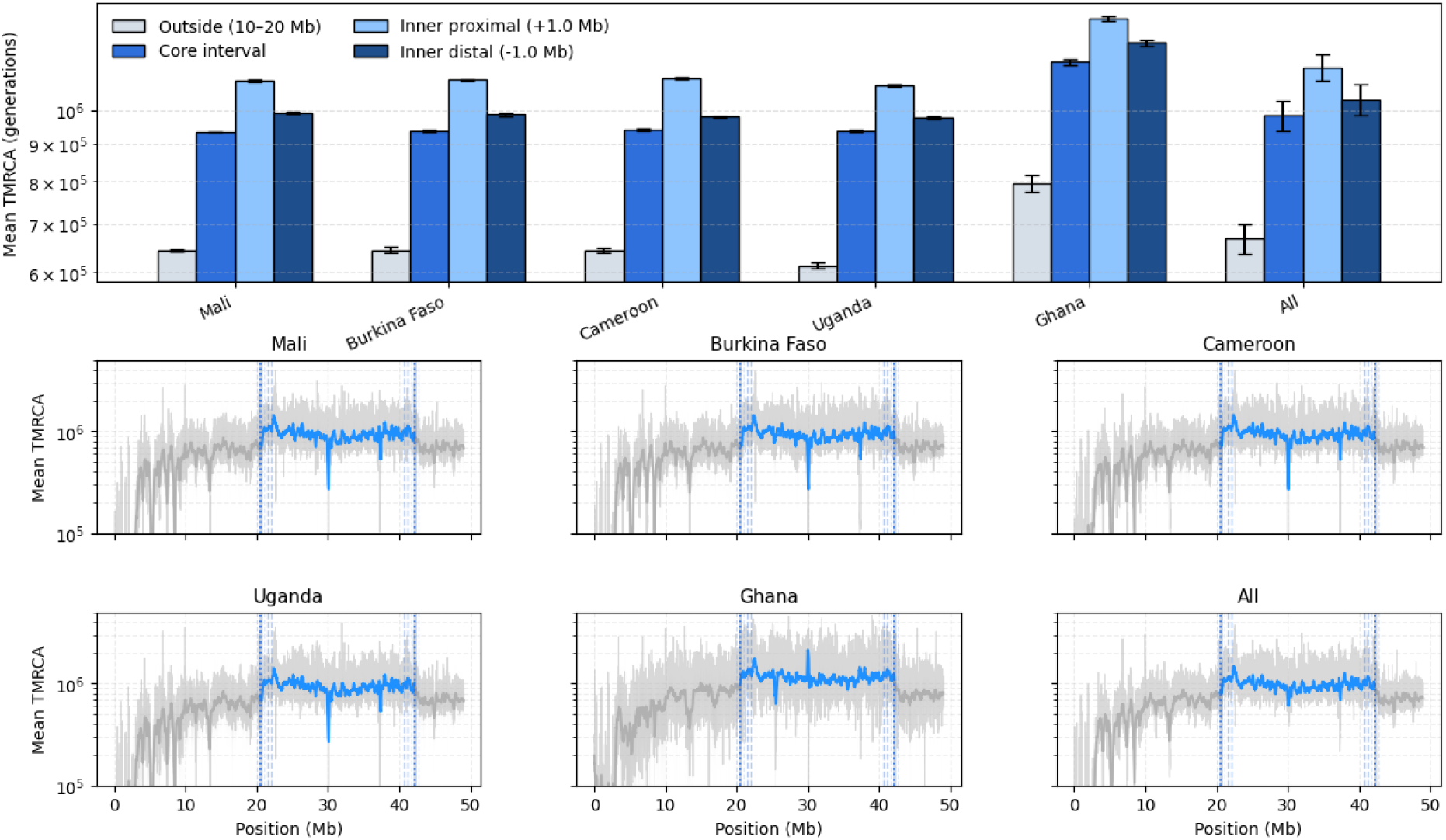
Coalescent-time structure across the *In(2L)a* inversion on chr2L. Top: mean TMRCA summaries for an outside background region (10–20 Mb), the full inversion core interval, and two interior 0.5 Mb windows positioned 1 Mb inside each breakpoint (mean ± s.e.m. across replicate trajectories; “All” aggregates populations). Bottom: genome-wide mean TMRCA curves shown per population and pooled (“All”), with vertical guides marking inversion boundaries, breakpoint-adjacent windows, and the interior sampling windows used for summary statistics.

To address these issues, we fine-tuned `cxt` on a filtered `stdpopsim` training set restricted to *N_e_* *>* 10^5^, reduced the window size ten-fold to 200 bp, and performed inference in 100 kb blocks. We additionally incorporated Ag1000G missing-data masks during fine-tuning to improve robustness to region-specific missingness.

Finally, because the Ghana population does not meet the 25-diploid sample size used elsewhere, we trained a small adapter model to handle reduced sample size, while jointly accounting for large N*_e_* and missing-data patterns.

The `Singer+Polegon` method was also run on the same samples. We tried two approaches to using `Singer+Polegon`— one with the “natural” parameterization of mutation and recombination rates, which for *Anopheles* yields a ratio of recombination:mutation `>>` 1, and a second where we held this ratio fixed at 1 (equivalent to the human parameterization). The former, “natural” parameterization yielded poor fits to observed diversity, particularly the marginal site frequency spectrum, thus the results presented here do not use mutation and recombination rates that reflect what is known about the *Anopheles* genome, although the resulting fits are superior with respect to matching observed diversity patterns.

Both `cxt` and `Singer+Polegon` successfully recover the pronounced reduction in TMRCA at *Rdl*. Importantly, the depth of this trough is not uniform across populations: some populations, such as Ghana, show a sharper, more localized depression, whereas Uganda shows no depression in TMRCA at all. This pattern parallels published differences in the frequency of *Rdl* insecticide-resistance alleles among these populations (Grau-Bové et al. 2020), supporting the interpretation that the inferred coalescent-time signal reflects geographically heterogeneous selection.

In comparing their outputs, a key difference is in the behavior in regions with missing data. `Singer+Polegon` consistently infers strong dips in the TMRCA in regions with missing data (see Figure S9), presumably as a result of miscalibrated expectations of mutation rate in those regions. However, because `cxt` is explicitly trained to deal with missing data, no such artifacts are inferred.

`SMC++` also detects a noticeable reduction in TMRCA around the *Rdl* sweep, but its blockwise representation limits how precisely this signal can be localized along the genome. Even after scaling the `SMC++` output to match the true genomic span of the region and applying manual calibration so that its genome-wide diversity patterns align with the `cxt` and Singer+Polegon panels, the inferred trough remains discretized and slightly shifted. This behavior is expected: `SMC++` models the genome using coarse blocks under a global demographic process, so narrow sweeps can be detected, but their boundaries are blurred by the block structure. As a result, `SMC++` can identify strong selective events, but it is less able to resolve their precise genomic breakpoints.

### Inversion interior coalescence signal and dating context

To characterize how the *In(2L)a* inversion shapes genealogical age across chr2L, we summarize mean TMRCA in (i) an outside background region (10–20 Mb), (ii) the full inversion core interval, and (iii) two 0.5 Mb windows sampled *inside* the inversion, offset by 1 Mb from each breakpoint to avoid the immediate transition zone where mapping artifacts, missingness, and recombination suppression can confound interpretation. Averaged across populations, coalescent times are elevated inside the inversion relative to the outside background: outside (10–20 Mb) = 6.676 × 10^5^ ± 7.148 × 10^4^ generations, core interval = 9.830 × 10^5^ ± 1.017 × 10^5^, inner proximal (+1.0 Mb) = 1.144 × 10^6^ ± 1.077 × 10^5^, and inner distal (−1.0 Mb) = 1.034 × 10^6^ ± 1.133 × 10^5^ (mean ± s.d. across populations). This ordering (outside < core < inversion interior) is consistent with long-lived haplotype structure maintained within the inversion, as expected under reduced effective recombination between arrangements and/or long-term maintenance of both karyotypes. In practice, focusing on the inversion *interior* provides a conservative summary that reflects the inversion’s long-term genealogical signal while reducing sensitivity to breakpoint-adjacent artifacts.

## Discussion

In this study, we cast population-genetic inference as a language-modeling problem by training a modified GPT-2 model to learn the sequential structure of the coalescent with recombination as a conditional stochastic process. In spirit, the method is aligned with SMC-based approaches such as `SMC++`, but it replaces the explicit HMM likelihood—and its enforced first-order dependence—with attention-based sequence modeling. Modeling assumptions and priors are supplied implicitly through simulation during training, rather than specified analytically at inference time.

The core of `cxt` is an autoregressive conditional generator: given observed mutations in a window, the model samples a discrete distribution over pairwise coalescence times and then extends predictions along the chromosome by conditioning on its own previously generated states. Repeating this procedure yields an approximate posterior over TMRCA trajectories that can be sampled efficiently in parallel. This approach differs fundamentally from prior deep learning methods such as `ReLERNN` (Adrion, Galloway, and Kern 2020) or `CoalNN` (Saada et al. 2023), which treat inference along the chromosome as a supervised regression task, and produce a single point estimate from sequence data. In contrast, `cxt` uses a language model to learn the underlying conditional stochastic process.

In this work, `cxt` produces genome-wide TMRCA estimates for a diploid individual in minutes on a single GPU, and it can alternatively focus on a specific region and decode many pairs locally. Comparisons to `SMC++` and `Singer+Polegon` show that `cxt` consistently outperforms `SMC++` in terms of accuracy of TMRCA inference (Table 1), and matches the state-of-the-art ARG-based method `Singer+Polegon`, a strong benchmark given its joint use of all samples. Achieving comparable performance without explicit ARG modeling highlights the efficiency of the language-model approach. These computational and statistical properties motivate application to empirical datasets spanning both canonical and challenging inference regimes.

To assess the performance of cxt on empirical data, we applied it to two empirical systems with very different mutation-to-recombination regimes: (i) a targeted validation in humans of the *HLA* and *LCT* regions of the genome— known coalescent time outliers, and (ii) a chromosome-scale scan of chr2L in Anopheles, highlighting the extensibility of our model to settings with huge population sizes and pervasive missing data that challenge standard statistical estimation.

In humans, the *LCT* locus provides a canonical example of recent, strong, and geographically structured selection. Lactase persistence is driven primarily by regulatory variation upstream of *LCT*, including variants located in intronic sequence of the neighboring *MCM6* gene that modulate enhancer/promoter activity (Swallow 2003; Tishkoff et al. 2007). Population-genetic analyses indicate that derived lactase-persistence haplotypes rose rapidly in frequency over roughly the past 5,000–10,000 years, consistent with gene–culture coevolution in the context of dairying (Bersaglieri et al. 2004; Itan et al. 2009). Our pairwise TMRCA estimates at *LCT* reflect this known history cleanly, recovering a pronounced local dip in coalescence times centered on the locus. In our sample, some pairs exhibit a TMRCA younger than 10 000 years, substantially younger than the average across the region, but simultaneously we note that there is extensive variation across pairs that likely reflects that LCT arose from standing variation (Peter, Huerta-Sanchez, and Nielsen 2012), and thus the locus harbors significant stochasticity in the underlying genealogies.

In the *HLA* region, we observe exceptionally ancient separation times among alleles. For several genes, pairwise TMRCAs exceed commonly cited human–chimpanzee divergence times, consistent with trans-species polymorphism under long-term balancing selection. The *HLA* locus is among the most polymorphic regions of the human genome, likely driven to extreme variation by arms-race dynamics at the immune genes it encodes. Accordingly, there is abundant evidence that some *HLA* alleles have been maintained for tens of millions of years by balancing selection (Hedrick 1998; Azevedo et al. 2015; Fortier and Pritchard 2025). In our sample of British individuals, the *HLA* region is a clear outlier in the chromosome-wide distribution of coalescence times (Figure 6, top right). Across multiple genes, we observe pairwise TMRCAs exceeding 10 million years. While surprisingly ancient, these estimates are nonetheless in agreement with recent population-genetic dating in (Deng, Song, and Nielsen 2025). Together, *LCT* and *HLA* provide complementary extremes, recent directional selection versus long-term balancing selection, allowing us to validate `cxt` across nearly the full dynamic range of human genealogical timescales, from thousands to tens of millions of years.

Unlike the human analyses, which primarily serve as validation against well-characterized loci, the Ag1000G application reveals previously unresolved spatial and temporal structure within ancient inversions and insecticide-resistance sweeps. These mosquito data also present a substantially more challenging setting for coalescent inference. In particular, the Ag1000G data combine very large effective population sizes, pervasive and spatially heterogeneous missingness, and uneven sample sizes across populations. Together, these factors challenge both diversity-based summaries and likelihood-based decoding, and provide a stringent test of whether `cxt` can be adapted to real data with non-ideal sampling and observation patterns.

We highlight chromosome 2L which carries a number of known targets of insecticide resistance as well as a well studied inversion, *In(2L)a*. Considering first the chromosomal landscape of estimates (Figure S9, bottom panels), we observe that the *In(2L)a* inversion region displays generally deeper coalescent times than the surrounding genomic background, with particular peaks in age near its breakpoints; consistent with prior observations of elevated diversity within this inversion (Love et al. 2016). Previous age estimates of the inversion using phylogenomics have aged the origin of the inversion to more than two million years ago (Fontaine et al. 2015), before the origin of the *An. gambiae* complex of species. We find that the regions proximal to the inversion breakpoints show ages on the order of ∼ 1.1 million generations, consistent with this deep origin. This pattern likely reflects a combination of suppressed recombination between inverted and standard arrangements, as well as long-term balancing selection maintaining both arrangements in the population. While the deep origin of *In(2L)a* is well established, how such an ancient inversion has been maintained within populations, and whether its evolutionary history is uniform across its length, remains less well understood. The pronounced and spatially structured coalescent age peaks near the inversion breakpoints indicate that different regions of the inversion have experienced markedly different genealogical histories since its origin. This suggests that recombination suppression and long-term maintenance have acted heterogeneously across the inversion. More broadly, it demonstrates that population-scale genealogical inference can reveal biologically meaningful structure in the evolutionary dynamics of ancient chromosomal inversions beyond a single global age estimate.

Along chromosome 2L, a significant outlier is the well-studied *Rdl* locus, which confers resistance to dieldrin and other cyclodiene insecticides (R. H. Ffrench-Constant et al. 1993; Thompson, Steichen, and R. Ffrench-Constant 1993; Du et al. 2005). Resistance to dieldrin appeared in Nigerian populations only 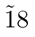 months after the initial onset of insecticide control efforts in 1954 (Elliott and Ramakrishna 1956) as a single, dominant allele (Davidson and Hamon 1962). Population genetic evidence points to a strong selective sweep at *Rdl* as well as a complex history of introgression that extends between species and inversion karyotypes (Grau-Bové et al. 2020). Our coalescent time estimates fill in this picture; we find clear evidence of recent selection at the locus, with short coalescent times among individuals in populations, such as Ghana, in which the resistant *Rdl* is at high frequency. Conversely in east African samples, where resistance is not found, coalescent times at the locus are indistinguishable from the local background.

Generally, our youngest time estimates at *Rdl* are on the order of hundreds to thousands of years, which would predate the introduction of broad insecticide use. Two things could be pushing back our dates here: i) it could be that the mutation was segregating at low frequency before the onset of selection, and ii) our estimates average over sites, and are not dating the origin of the *A296G* nonsynonymous mutation that confers resistance in *An. gambiae*.

Taken together, these applications show that `cxt` remains effective on large empirical datasets, including local genomic regions that deviate substantially from the evolutionary scenarios represented in its simulation-based pretraining, which acts as an implicit prior, such as targets of selective sweeps and long-term balancing selection. The Ag1000G analyses, in particular, highlight a practical advantage of simulation-based models: the ability to adapt to new evolutionary regimes with minimal targeted modification. By fine-tuning on high–*N_e_* simulations, reducing the window size to 200 bp, incorporating empirical missing-data masks, and adding a lightweight adapter for small sample sizes, `cxt` remained accurate in settings where traditional ARG MCMC and SMC-based methods face computational or modeling limitations. This flexibility, combined with amortized inference, positions `cxt` as a practical and scalable tool for population-genomics studies as community resources such as `stdpopsim` continue to expand to encompass additional species, demographic histories, and recombination landscapes.

In summary, we introduce a language-model-based inference engine designed to scale with the growing `stdpopsim` catalog. By directly learning a conditional stochastic process, `cxt` absorbs prior knowledge directly through simulation-based training, without requiring new analytic likelihood derivations for each scenario. As a result, `cxt` is a fast and accurate tool for TMRCA inference, and provides a flexible framework that can be adapted to novel evolutionary settings.

## Methods

We start by providing a detailed overview of the methods introduced in our study. First, we briefly introduce the concept of **next-coalescence prediction**. Second, we describe the parameterization of coalescent simulation and the way we formatted mutational data in order to present it to our network. Third, we outline the architecture of the network itself, which is derived from the GPT-2 decoder-only model. Finally, we describe the various experiments conducted to benchmark our method according to the objectives described above.

### Next-coalescence Prediction

We introduce as our prediction task, *next-coalescence* prediction, which we define to mean the next point in time in which a focal (or pivot) pair of sequences find common ancestry as we move across (say from left to right) a sequence. Our method processes 500 genome windows (most commenly e.g. 500 × 2 Kb or 1Mb; or 500 × 200 bp or 0.1 Mb) of sequence. For the purposes of this paper we use a sample size of 25 diploid individuals, but this is arbitrary and we provide a method to train a light-weight adapter generalizing to required input sizes (or recommend sub-sampling and averaging for larger sample sizes). Our model focuses on a pivot pair of chromosomes, also non-as distinguished pairs in `SMC++` terminology, from which we predict pairwise coalescence events, or more specifically a probability distribution from which the next coalescence event can be sampled for that pivot pair along the sequence for each of the 500 windows. This describes an iterative process by which the predicted coalescence time is concatenated with the mutational context, forming the next input to the model to predict the next coalescence event (see Figure 1). We refer to this procedure as *next-coalescence prediction*. We continue by describing the processing of the genotype matrix as obtained from coalescent simulation before presenting the architecture and listing the specific simulation parameterization next.

### Processing of Coalescent Simulations

We begin with a genotype matrix *G* ∈ ℤ*^N×M^* containing *N* diploid samples across *M* sites, from coalescent simulations (here `msprime`). Phasing is not required for real data, as using both haplotypes per individual yields the same pivot–pair conditioning used throughout.

A high-level description is provided here, with an explicit version in Algorithm 1 and a schematic overview in Figure S1. The core preprocessing consists of extracting mutation patterns for each pivot pair using logical operations—-heterogeneous differences via `XOR` and homogeneous matches via `XNOR`—-and then weights these these pivot-pair vectors by their site-frequency spectrum (SFS) counts computed across all remaining samples. This converts the otherwise binary mutation indicators from the logical opperations into coarse age-informed signals, ensuring that the inferred pairwise coalescence times are driven by the allele frequency spectrum present in the entire sample.

This procedure is applied in sliding windows of increasing size (4–128 K b) with a 2 K b stride, which allows for rescaling when inference is performed at different resolutions.

In summary, the model input consists solely of the genotype matrix and variant positions; from these, we derive windowed, SFS-weighted mutation patterns for each pivot pair, which the language model uses to estimate average coalescence times per window. These inputs arise naturally from `msprime` simulations or from empirical genotype data such as VCFs.

### Implementation of the Decoder-Only Architecture

We implement a decoder-only transformer closely aligned with GPT-2 (Radford et al. 2019), rather than the original encoder–decoder formulation (Vaswani et al. 2017), and adapt it to the structure of population genetic data. In NLP, text is tokenized and mapped to embeddings via a learned lookup table; in contrast, the mutation patterns in genomic windows form a vast, effectively continuous space that makes discretization impractical. Instead of defining a bespoke tokenizer, we directly project SFS-weighted mutation patterns into the latent space using a fully connected embedding module (Algorithm 2), which accepts tensors of shape *B* × *Z* × *W S* × *K* × *N* and outputs *B* × *K* × *E*. Here, *B* is batch size, *Z* the logical vector distinguishing heterozygous and homozygous states, *W S* the number of window sizes, *K* the number of windows, *N* the number of samples, and *E* the embedding dimension.

Coalescence times are represented by partitioning log-scaled values into 324; Table S1) discrete intervals, each mapped through a small embedding table. The model then concatenates mutation and coalescence embeddings so that every context window carries both the local mutational signal and the discretized temporal state. This setup allows the same model to perform inference across diverse evolutionary scenarios without re-training, since the inputs already encode the relevant population-genetic context.

To encode genomic position, we adopt rotary positional embeddings (Su et al. 2023), which efficiently capture spatial structure and require no adjustment under fixed-stride windows (e.g., 2 kb). The resulting sequence fed to the transformer consists of 500 mutation embeddings, a start token, and the coalescence embeddings, yielding a fixed latent length of 1001; padding exists but is unused. A dropout rate of **0.1** is applied for regularization. Both narrow (single-scenario) and broad (multi-scenario) models (`cxt`-narrow and `cxt`-broad respectively) share this architecture, with parameterization summarized in Table S1. Further implementation details are available at https://github.com/kr-colab/cxt.

### Sample-size and accessibility adaptation

`cxt` models are trained at a fixed sample size, which determines the number of SFS channels used to represent the mutational context. Empirical datasets may contain fewer samples than the pretrained model (e.g. the Ghana population in Ag1000G), and retraining the full model for each sample size would be inefficient. To enable transfer to smaller cohorts, we introduce a lightweight input-adaptation stage that operates before the pretrained decoder-only transformer.

#### Sample-size adapter

The adapter maps SFS features computed from a reduced cohort to the dimensionality expected by the pretrained model. After this transformation, data are processed by the original projection layers and transformer without further modification. In this way, the adapter modifies only how mutational information is presented to the model, while preserving the pretrained representation of the coalescent process.

The adapter is implemented as a small residual neural network applied independently to each genomic window. It consists of layer normalization followed by a gated bottleneck that rescales and recombines SFS channels derived from the smaller cohort. A residual connection projects the input directly to the target dimensionality, and a learnable scaling parameter initialized near zero controls the magnitude of the adapter’s contribution. As a result, the adapter behaves close to an identity mapping at the start of training and fine-tunes in a stable manner.

#### Training procedure

Adapter training is performed by freezing all parameters of the pretrained decoder-only transformer and updating only the adapter parameters. The training objective is identical to that of the full model, namely prediction of the local pairwise coalescence time in each genomic window. Because the adapter contains far fewer parameters than the backbone, it converges rapidly and can be trained in a small number of epochs using simulations generated at the target sample size.

To accommodate small but systematic shifts in input statistics, we additionally unfreeze the LayerNorm parameters and the final transformer blocks and train them at a substantially reduced learning rate. This restricted fine-tuning improves stability when transferring to smaller cohorts while avoiding degradation of the pretrained coalescent prior.

#### Fine-tuning to genomic inaccessibility

Empirical genomes contain extended regions that are systematically uncallable, producing structured missingness patterns that differ from idealized simulations. To account for this, we fine-tune the adapted model on training examples in which genomic inaccessibility is imposed explicitly using an empirical accessibility mask.

For each simulated tree sequence, we select a contiguous genomic segment and delete intervals corresponding to unaccessible sites from the tree sequence, ensuring that both mutational and genealogical information reflect the empirical observation process. In parallel, we compute the fraction of unaccessible bases per window across multiple scales and inject this signal into a dedicated input channel, providing the model with explicit information about local callability. This fine-tuning step uses the same training objective as above and updates the same restricted set of parameters (adapter, LayerNorms, and final transformer blocks), allowing the model to retain its pretrained coalescent representation while calibrating to irregular observation patterns in real genomes.

### Calibration of the Molecular Clock

Although `cxt` is mutation-rate agnostic during training, its raw TMRCA predictions must be placed on an absolute timescale for demographic interpretation. We therefore apply a simple calibration step that rescales predicted TMRCAs so that the expected nucleotide diversity matches the observed diversity across the full context window. This correction is typically negligible when the data resemble the training scenarios, but becomes useful under atypical mutation rates or demographic histories. To avoid collapsing variability across replicates, we treat the correction factor probabilistically rather than deterministically. A full derivation, including the Poisson–Gamma model used to sample per-replicate correction factors, is provided in Supplementary Materials .

### Estimating Population Size from Coalescence Rates

To convert predicted TMRCAs into population size trajectories, we estimate the instantaneous coalescence rate on a logarithmic time grid and invert it to obtain *N_e_*(*t*) = 1/(2*λ*(*t*)). This provides a nonparametric demographic reconstruction directly from the model’s TMRCA samples. Details, including windowing, discretization, and treatment of the tail interval, appear in Appendix .

### Datasets and Training Description

The precise coalescence parameter initiations used for training of both models are listed in the supplementary material, organized into distinct sets, a single parameter set for the narrow model, roughly approximating a constant demography human scenario, and the broad model extensions including dataset encompassing most of stdpopsim v0.2 and v0.3, the latter used explicitly for validation not training.

We implemented our models using PyTorch Lightning to streamline code management, including multi-GPU training and loss tracking. Matrix multiplication precision was set to float32 (medium), relying on PyTorch Lightning’s internal precision handling, and we did not use additional manual autocasting with ’bf16-mixed’ during training. To optimize memory efficiency and storage, datasets were stored in float16 format.

The model was trained for five epochs using the AdamW optimizer with a learning rate of 3 × 10*^−^*^4^, which was gradually decreased during training via cosine annealing.

Fine-tuned models were trained for only two epochs on their respective datasets, using a learning rate reduced by a factor of ten. For more information about specific training commands, we refer to the online manual.

Each training sample of a batch consists of 500 windows, followed by a start token, which is the second row of the embedding table for time discretization (the first row is reserved for potential padding, but is not used in this implementation). This is then followed by 500 coalescent times. The target sequence is simply a right-shifted version of the coalescent times, beginning with the first coalescent event. Accordingly, the model is trained in the same manner as standard language models, using negative log-likelihood loss and a causal mask. The causal mask has been adapted for translation within the relevant attention layers. Specifically, we used a fused-causal attention mask, which lets the decoder distinguish mutation from coalescence tokens when “looking back” over the sequence: when predicting a mutation, the model can attend to all previous tokens, but when predicting a coalescence, it can attend to all earlier mutations and only to past (not future) coalescences. This preserves strict causal ordering for sequence decoding while keeping the full, fixed mutation history visible at every step.

### Environmental considerations

Although GPUs typically draw more instantaneous power than CPUs (e.g., 250–400 W for an NVIDIA A100 versus 225 W for a high-end server CPU such as the AMD EPYC 7742), they can substantially reduce wall-clock time for inference workloads (distinct from training, which requires on the order of 10 h on three A100s) that map well to highly parallel operations. This is particularly relevant for our amortized inference setting, where recombination-rate variation does not affect runtime, unlike in full-likelihood methods such as Singer. In practice, GPU acceleration can markedly shorten runtime in certain parameter regimes–especially large Ne-scaled scenarios—-so that even with a higher power draw, the total energy use and *CO*_2_ emissions per completed inference may be lower than on CPUs, simply because the task finishes much faster.

## Acknowledgments

We thank Scott Small for his assistance with processing the Anopheles data. We thank members of the Kern-Ralph co-lab for helpful discussions and feedback. We also thank the authors of `SMC++` and `Singer` for their implementations and for providing their code. Finally, we thank Negar Rahnamae for designing the logo for the `cxt` method. This work was supported by NIH awards R35148253 and R01HG010774.

## Data and Code Availability

Simulation code, model training workflows, and accompanying documentation are available under a CC-BY-NC 4.0 license at https://github.com/kr-colab/cxt, with additional resources at https://cxt.readthedocs.io/en/latest/. Publicly available data from the 1000 Genomes Project were used for evaluation.

## Supplementary Material

### Things we tried as well

#### Preprocessing mutational patterns

We explored representing local genealogical configurations via a discrete “codebook” of mutation patterns, allowing the model to operate on higher-level coalescent motifs rather than raw mutation arrays. In practice, the combinatorial growth with sample size and window length made codebook learning brittle, non-extensible, and poorly suited for scenarios requiring inference of multiple coalescent times. Performance degraded sharply outside the specific training regimes.

#### Hand-rolled convolutions via multi-scale windows

To mimic convolutional behavior, the cur-rent model uses multiple window sizes in the embedding stack. Single-scale tokenization consistently undercaptured rare, localized events such as sharp TMRCA dips. Although we considered adding true convolution operators, the multi-scale approach remained the simplest, at the cost of increased storage and additional input processing requirements.

#### Architectural upgrades with limited impact

We evaluated more advanced attention variants (linear attention, multi-query attention) but observed negligible gains given our short contexts (about 500 tokens per 1 Mb). These optimizations mainly benefit very long-sequence models, and the added engineering complexity was not justified.

#### Splitting the encoder and decoder

We also considered a cleaner encoder-decoder separation for improved maintainability and the possibility of halving context length by compressing the mutation stream prior to temporal decoding. While promising-especially in light of simpler key-value caching-the monolithic GPT-like architecture appeared conceptually simpler, though we may return to this design in future models.

#### Training mixtures without repetition

Early training experiments drew demographies randomly without repetition across epochs. This prevented the model from seeing the same statistical structure often enough to form stable internal representations and led to degraded inference accuracy. Reintroducing repetition-training many iterations of the same demography with different random seeds-proved essential for stable learning and effective generalization.

### Algorithmic Details of Preprocessing

For processing the input the complete algorithmic details are provided in Algorithm 1 below. This describes the internal construction of the tensor to be ingested by our model.

The input module projecting SFS values per window of specified size into latent space in parallel for 500 windows, 4 window sizes, and 2 states (heterogeneous vs. homogeneous) and B batches.

Hyperparameter specifications of our narrow and broad model.

**Table S1:** Token-Free Decoder Model Configuration.

| Parameter | Narrow Model | Broad Model |
| --- | --- | --- |
| Number of Layers ($n_{\text{layer}}$) | 6 | 10 |
| Number of Attention Heads ($n_{\text{head}}$) | 4 | 4 |
| Embedding Dimension ($n_{\text{embd}}$) | 400 | 400 |
| Dropout Rate | 0.1 | 0.1 |
| Bias in Linear Layers | False | False |
| Number of Samples per Forward Pass | 50 | 50 |
| Sample Scale Embedding Factor | 2 | 2 |
| Output Dimension ($\text{output}_{\text{dim}}$) | 326 | 326 |
| Combined Feature Dimension ($\text{combined}_{\text{dim}}$) | 1001 | 1001 |
| **Total parameters** in millions | $\approx 11.7$ | $\approx 19.5$ |

The following sections contains detailed information about dataset parameter initiation and dataset sizes:

#### Algorithm 1

ProcessGenotypeMatrixAndPositions (Vectorized)

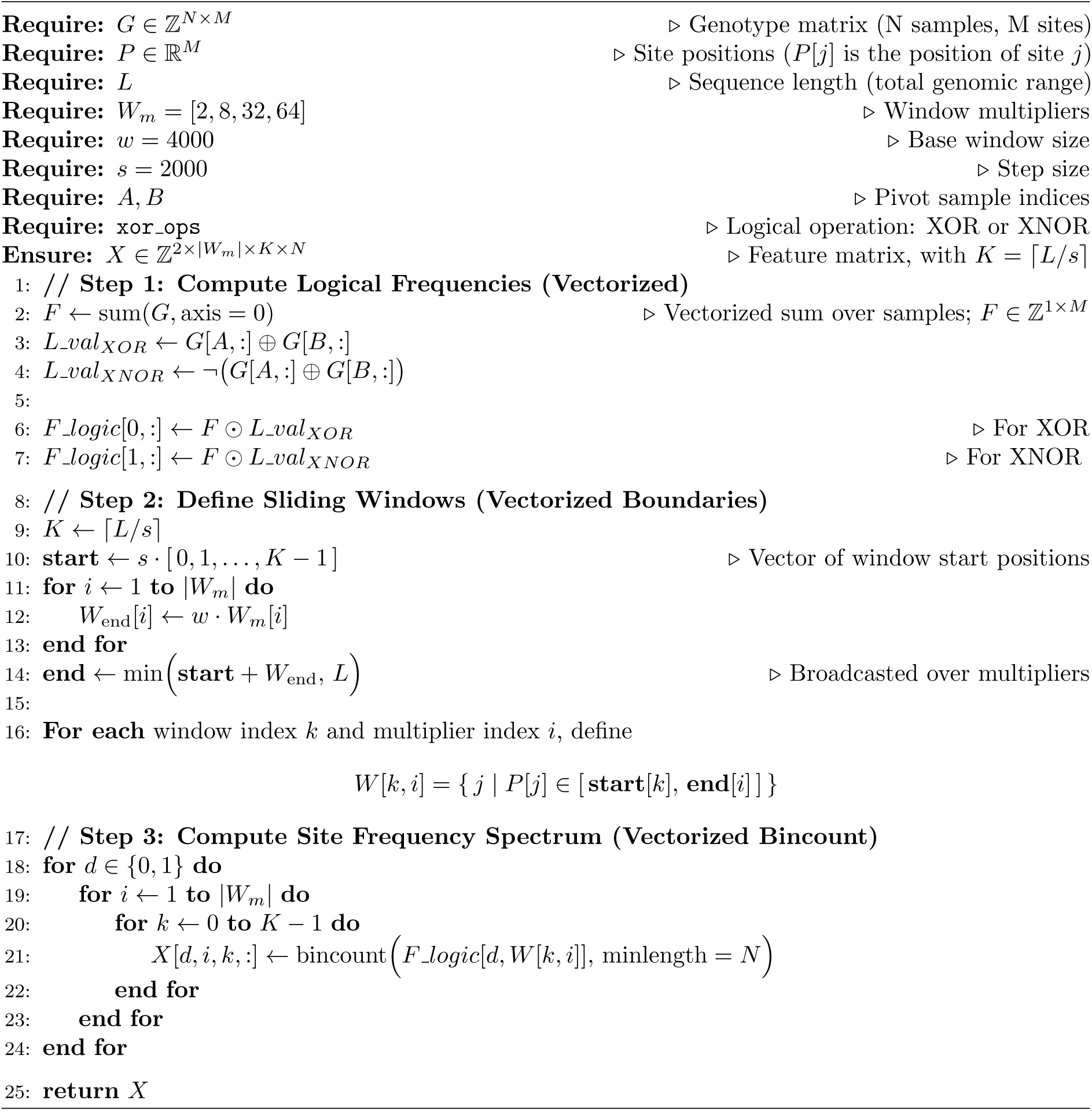

#### Algorithm 2

ProjectionReplacingEmbeddingTable

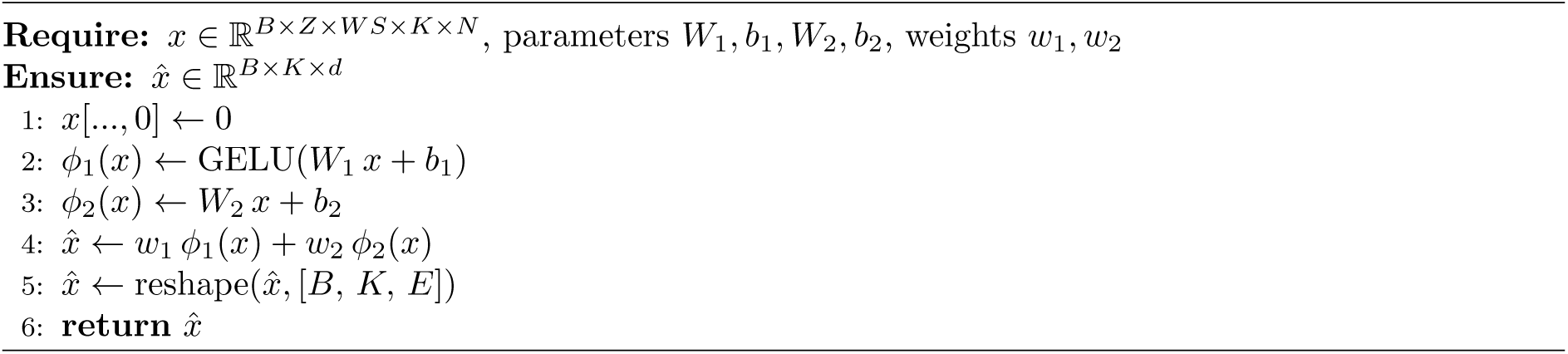

We organized our training data into four distinct sets. The first is a large base dataset, used to train the narrow model, with parameters chosen to approximate human simulations. This set consists of a single demographic scenario with constant population size (see Table S2). We also performed fine tuning of the narrow model for a few auxiliary models (Table S3). The third dataset is the stdpopsim v0.2 dataset (see Table S4), the main dataset used in this study that we call the broad model. The final dataset comprises stdpopsim v0.3 models, which are used for evaluation (see Table S5).

### Calibration of the Molecular Clock

`cxt` output is run in replicates (default 15), that together provide a distribution over TMRCA trajectories. Although the language model is agnostic to mutation rate (in the sense that this is not provided as an explicit input, and the model sees data generated under many mutation rates during training), it will generally be desirable to condition on a particular mutation rate/molecular clock for N*_e_* calibration purposes. This is done by scaling the raw predictions such that expected nucleotide diversity matches the observed nucleotide diversity across the full context (e.g. the 1Mb sequence). Empirically, this first-order correction is most helpful when the model is applied to data that is atypical relative to the training simulations (see Figures S2 and S3). Crucially, the relative TMRCA is highly accurate even in out-of-distribution scenarios, so that the first order correction is generally sufficient to make bias and mean squared error constant across a wide range of scenarios. When the input data are from a demographic scenario used in the training simulations, the impact of this correction becomes negligible (Figure **??**, compared to Figure 2).

One issue is that a deterministic first-order correction will reduce variability in `cxt`’s “posterior” samples– for example, TMRCA trajectories that are constant across the windows but have distinct values across replicates, would be all collapsed to the same value. Thus, we model the correction factor probabilistically to preserve replicate variability. For a given pivot pair *i*, we assume the observed mutation count `y`*_i_* follows a Poisson model:

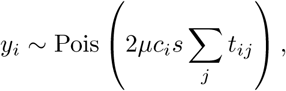

where *s* is the window size, *µ* is the mutation rate, t*_ij_* is the predicted TMRCA in window *j* of replicate *i*, and *c_i_* is the per-replicate correction factor. Under an improper constant prior, this yields a posterior:

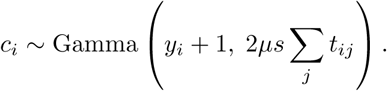

We then sample an independent *c_i_* for each replicate and apply it to all windows in that replicate. This propagates uncertainty from diversity estimation into the corrected predictions. In practice, the difference is negligible when mutation counts are large, as the posterior concentrates around the maximum likelihood estimate. However, when mutation counts are sparse or when the model produces flat TMRCA profiles, this stochastic correction helps avoid degeneracies (e.g., where there are no mutations and the MLE is undefined).

### Estimating Population Size via Instantaneous Inverse Coalescence Rates

We estimate the instantaneous coalescence rate from a set of ancestor coalescence times by discretizing time into logarithmically spaced windows (*T_max_* = 40). Specifically, we define the time grid using

**Table S2:** Base Dataset (Approx. for Human Parameters)

| Parameter | Value |
| --- | --- |
| Population Model | Constant Demography |
| Population Size | $2 \times 10^4$ |
| Mutation Rate | $1.29 \times 10^{-8}$ per Base |
| Recombination Rate | $1.28 \times 10^{-8}$ per Base |
| Simulations | $2 \times 10^6$ |
| Sequence Length | 1 Mb |

**Table S3:**
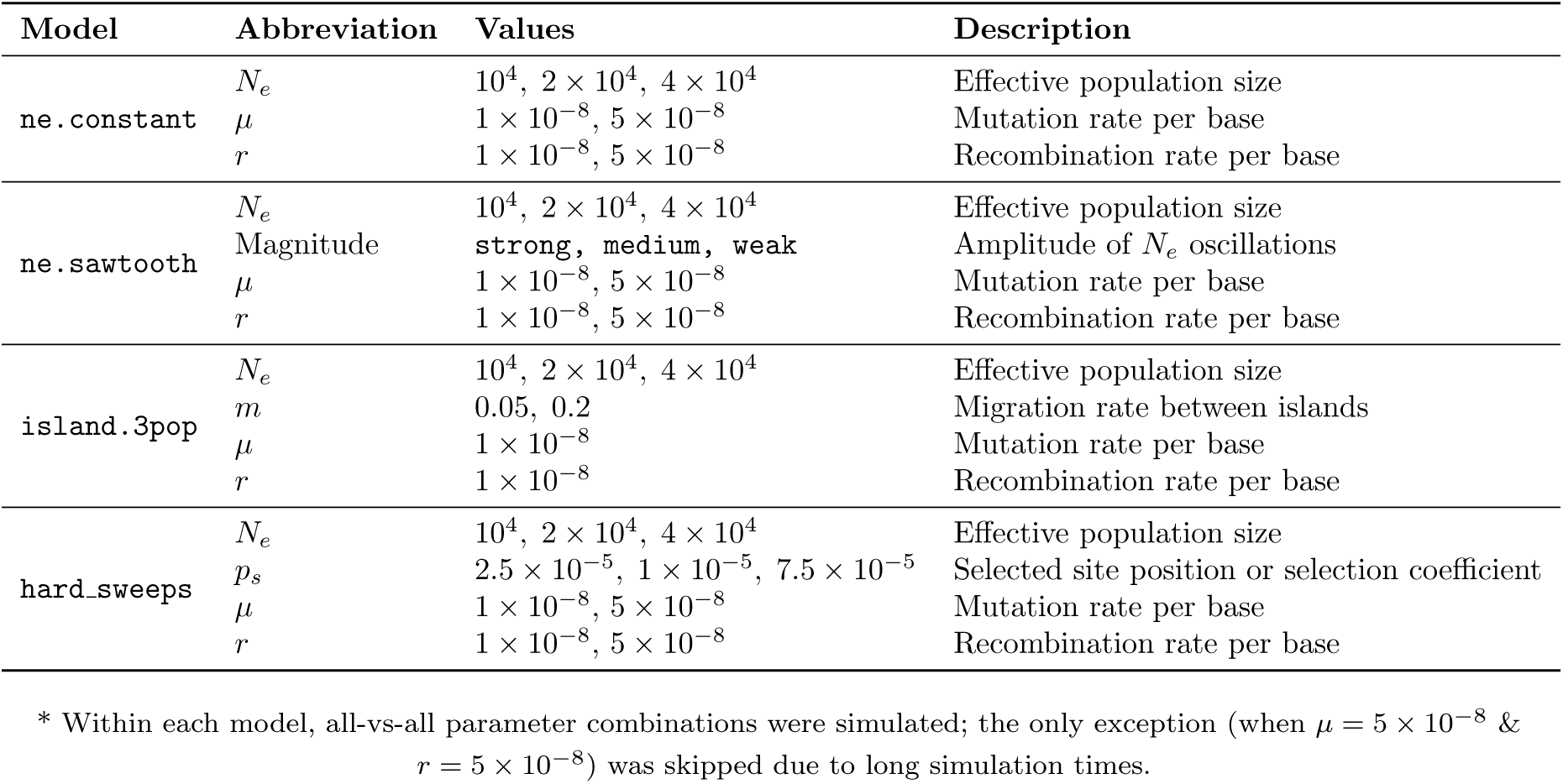
Base Dataset Extension (auxiliary Dataset): Model Parameters *.

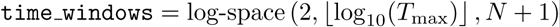

where *T_max_* is the maximum simulation time and *N* is the number of desired windows. We then explicitly set the first window edge to 0 by assigning `time windows[0] = 0.0`. This creates *N* non-overlapping intervals [*t*_0_, *t*_1_), [*t*_1_, *t*_2_), . . ., [*t_N−_*_1_, *t_N_* ) that cover the relevant timescale on a logarithmic scale, providing finer resolution for more recent coalescent events.

Let *T*_1_, *T*_2_, . . ., *T_n_* be the observed coalescence times. For each window [*t_i_*, *t_i_*_+1_), we compute the proportion of events falling into that bin, giving a discretized probability density function (PDF), denoted p*_i_*. The corresponding cumulative distribution function (CDF) is *F*(*t_i_*) = Σ*_j<i_ p_j_*. Assuming a memoryless coalescence process (i.e., a Poisson process), the instantaneous rate λ(*t*) satisfies:

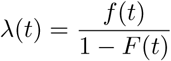

where *f*(*t*) is the density and 1 − *F*(*t*) is the survival function. Integrating this rate over the interval [*t_i_*, *t_i_*_+1_) gives:

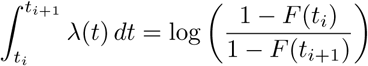

and dividing by the window width Δ*t* = *t_i_*_+1_ − *t_i_*, we estimate the average coalescence rate in the interval as:

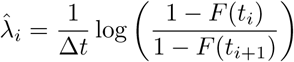

For the final window (where the survival function becomes too small), we estimate the rate via the inverse of the mean residual coalescence time:

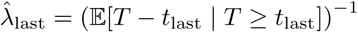

This approach provides a nonparametric, smoothed estimate of the coalescence rate as a function of time, suitable for demographic inference.

**Table S4:**
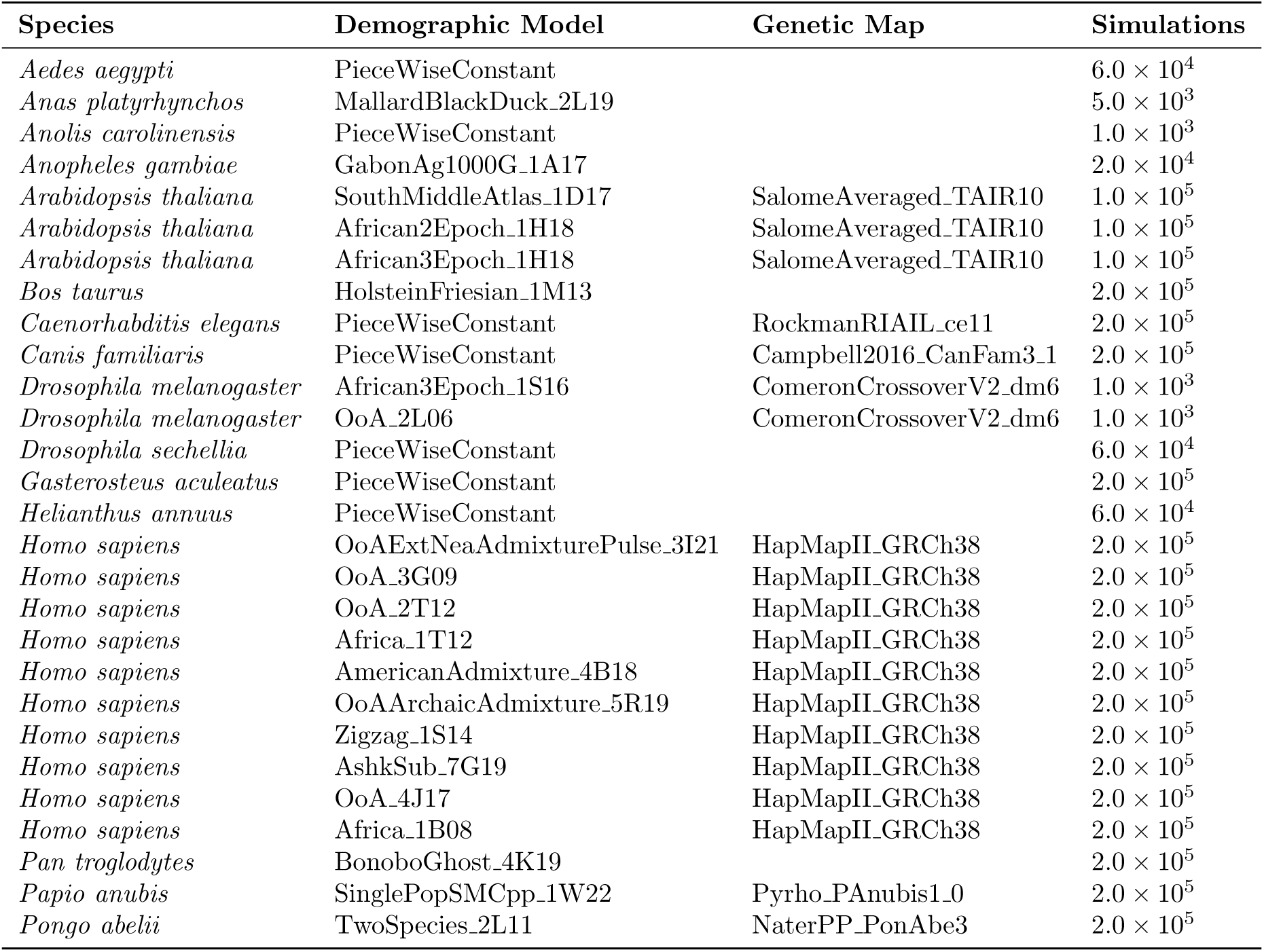
stdpopsim dataset (v0.2): Simulations configuration for training.

| Species | Demographic Model | Genetic Map | Simulations |
| --- | --- | --- | --- |
| *Aedes aegypti* | PieceWiseConstant | | $6.0 \times 10^4$ |
| *Anas platyrhynchos* | MallardBlackDuck_2L19 | | $5.0 \times 10^3$ |
| *Anolis carolinensis* | PieceWiseConstant | | $1.0 \times 10^3$ |
| *Anopheles gambiae* | GabonAg1000G_1A17 | | $2.0 \times 10^4$ |
| *Arabidopsis thaliana* | SouthMiddleAtlas_1D17 | SalomeAveraged_TAIR10 | $1.0 \times 10^5$ |
| *Arabidopsis thaliana* | African2Epoch_1H18 | SalomeAveraged_TAIR10 | $1.0 \times 10^5$ |
| *Arabidopsis thaliana* | African3Epoch_1H18 | SalomeAveraged_TAIR10 | $1.0 \times 10^5$ |
| *Bos taurus* | HolsteinFriesian_1M13 | | $2.0 \times 10^5$ |
| *Caenorhabditis elegans* | PieceWiseConstant | RockmanRIAIL_ce11 | $2.0 \times 10^5$ |
| *Canis familiaris* | PieceWiseConstant | Campbell2016.CanFam3.1 | $2.0 \times 10^5$ |
| *Drosophila melanogaster* | African3Epoch_1S16 | ComeronCrossoverV2_dm6 | $1.0 \times 10^3$ |
| *Drosophila melanogaster* | OoA_2L06 | ComeronCrossoverV2_dm6 | $1.0 \times 10^3$ |
| *Drosophila sechellia* | PieceWiseConstant | | $6.0 \times 10^4$ |
| *Gasterosteus aculeatus* | PieceWiseConstant | | $2.0 \times 10^5$ |
| *Helianthus annuus* | PieceWiseConstant | | $6.0 \times 10^4$ |
| *Homo sapiens* | OoAExtNeaAdmixturePulse_3I21 | HapMapII_GRCh38 | $2.0 \times 10^5$ |
| *Homo sapiens* | OoA_3G09 | HapMapII_GRCh38 | $2.0 \times 10^5$ |
| *Homo sapiens* | OoA_2T12 | HapMapII_GRCh38 | $2.0 \times 10^5$ |
| *Homo sapiens* | Africa_1T12 | HapMapII_GRCh38 | $2.0 \times 10^5$ |
| *Homo sapiens* | AmericanAdmixture_4B18 | HapMapII_GRCh38 | $2.0 \times 10^5$ |
| *Homo sapiens* | OoAArchaicAdmixture_5R19 | HapMapII_GRCh38 | $2.0 \times 10^5$ |
| *Homo sapiens* | Zigzag_1S14 | HapMapII_GRCh38 | $2.0 \times 10^5$ |
| *Homo sapiens* | AshkSub_7G19 | HapMapII_GRCh38 | $2.0 \times 10^5$ |
| *Homo sapiens* | OoA_4J17 | HapMapII_GRCh38 | $2.0 \times 10^5$ |
| *Homo sapiens* | Africa_1B08 | HapMapII_GRCh38 | $2.0 \times 10^5$ |
| *Pan troglodytes* | BonoboGhost_4K19 | | $2.0 \times 10^5$ |
| *Papio anubis* | SinglePopSMCpp_1W22 | Pyrho_PAnubis1_0 | $2.0 \times 10^5$ |
| *Pongo abelii* | TwoSpecies_2L11 | NaterPP_PonAbe3 | $2.0 \times 10^5$ |

**Table S5:** stdpopsim dataset (v0.3): Simulations configuration for evaluation and retraining.

| Species | Demographic Model | Simulations (for retraining) |
| --- | --- | --- |
| *Gorilla gorilla* | GorillaGhost_5P23 | $2.0 \times 10^5$ |
| *Oryza sativa* | BottleneckMigration_3C07 | $2.0 \times 10^5$ |
| *Rattus norvegicus* | PieceWiseConstant | $2.0 \times 10^5$ |
| *Sus scrofa* | PieceWiseConstant | $2.0 \times 10^5$ |

**Figure S1:**
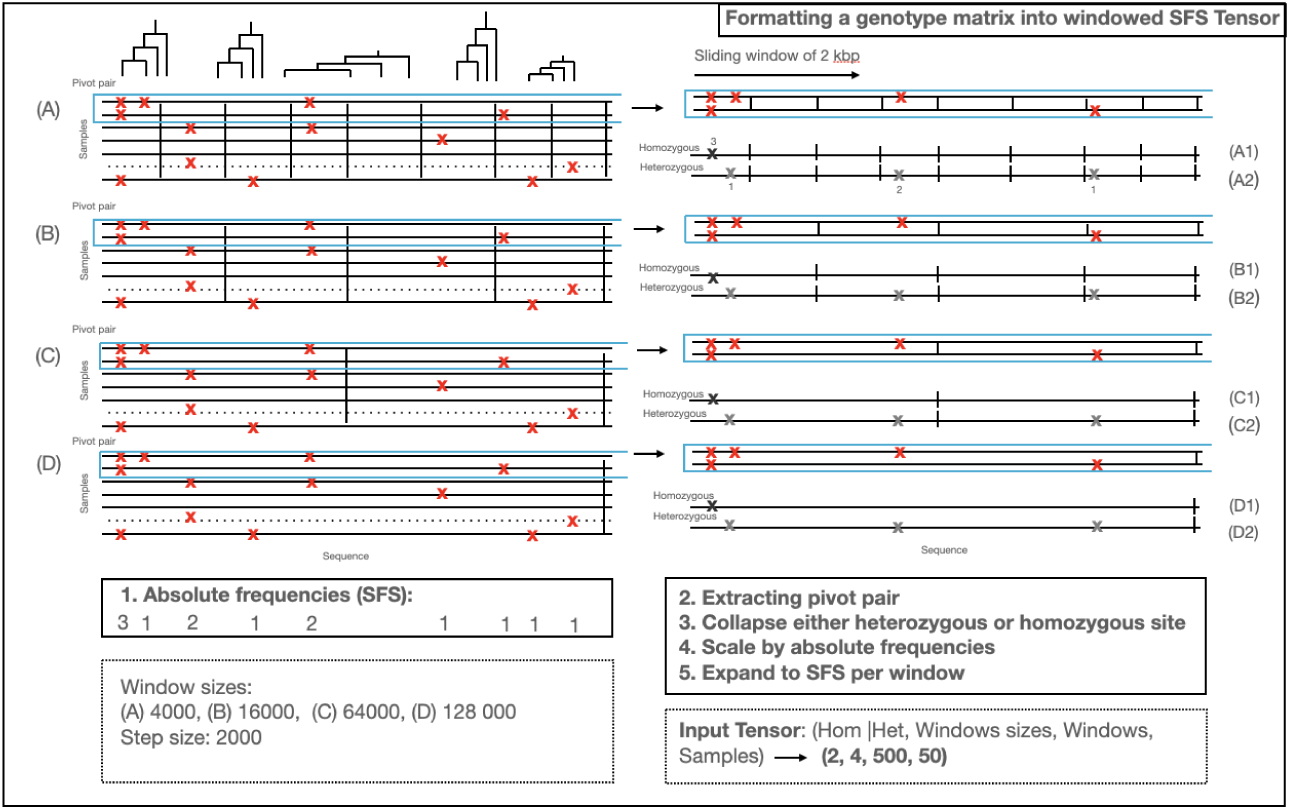
Processing of a phased genotype matrix, as detailed in Algorithm 1. This figure provides a schematic overview. Starting at (**A**), we compute absolute variant frequencies across 50 samples (a fixed hyperparameter) (1). A one-megabase region is divided into four-kilobase windows with a 2 Kb slide (not shown). We focus on a pivot pair (framed in blue), extract it, and classify sites as heterogeneous or homogeneous. Each site is scaled by its SFS class, and each window is expanded into the full sparse SFS dimensions of 50 (see input tensor, bottom right). This process repeats for three additional windows (**B–D**), yielding a complete input tensor of shape (2, 4, 500, 50) per pair, with the SFS aggregating information across all samples.

**Figure S2:**
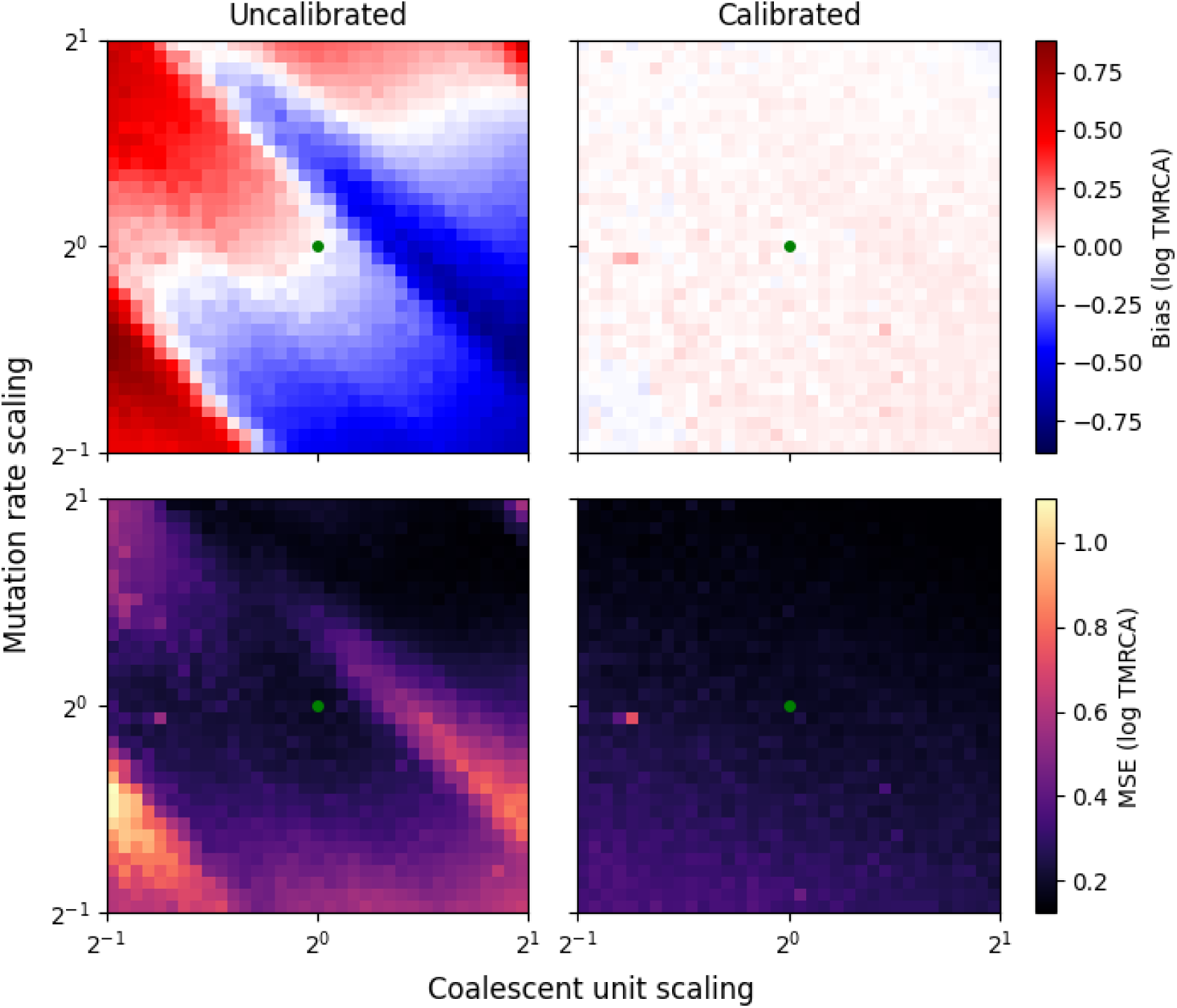
The impact of calibrating the molecular clock using observed nucleotide diversity (see section “Calibration of Molecular Clock” in methods), using simulations from perturbations of the Zigzag model in the stdpopsim catalog. The x-axis is a scaling of the Markov generator for the coalescent process, that increases the average TMRCA while keeping the shape of the site frequency spectra constant. The y-axis is a scaling of the mutation rate. The two columns correspond to uncalibrated and calibrated cxt predictions (e.g. in the former, the language model is predicting absolute TMRCA; and in the latter relative TMRCA). The two rows are bias (top) and mean squared error (bottom) relative to the true TMRCA, calculated from 30 independent simulations per grid cell. The unperturbed demography (which the language model sees during training) is the green point in the center of each heatmap. As the model is perturbed, the uncalibrated cxt predictions become substantially biased in a fashion that is nonlinear with regards to the magnitude of the perturbations (left column); reflecting the fact that the TMRCAs, mutation rate, and recombination rate are not jointly identifiable. However, relative changes in TMRCA are predicted accurately, so that centering the TMRCA trajectories around the observed nucleotide diversity for each pivot pair (e.g. conditioning on the mutation rate) removes the bias regardless of the degree of perturbation. Thus, after calibration (right column) the MSE for cxt predictions follows a gradient of mutational density (see Supplementary Figure S3).

**Figure S3:**
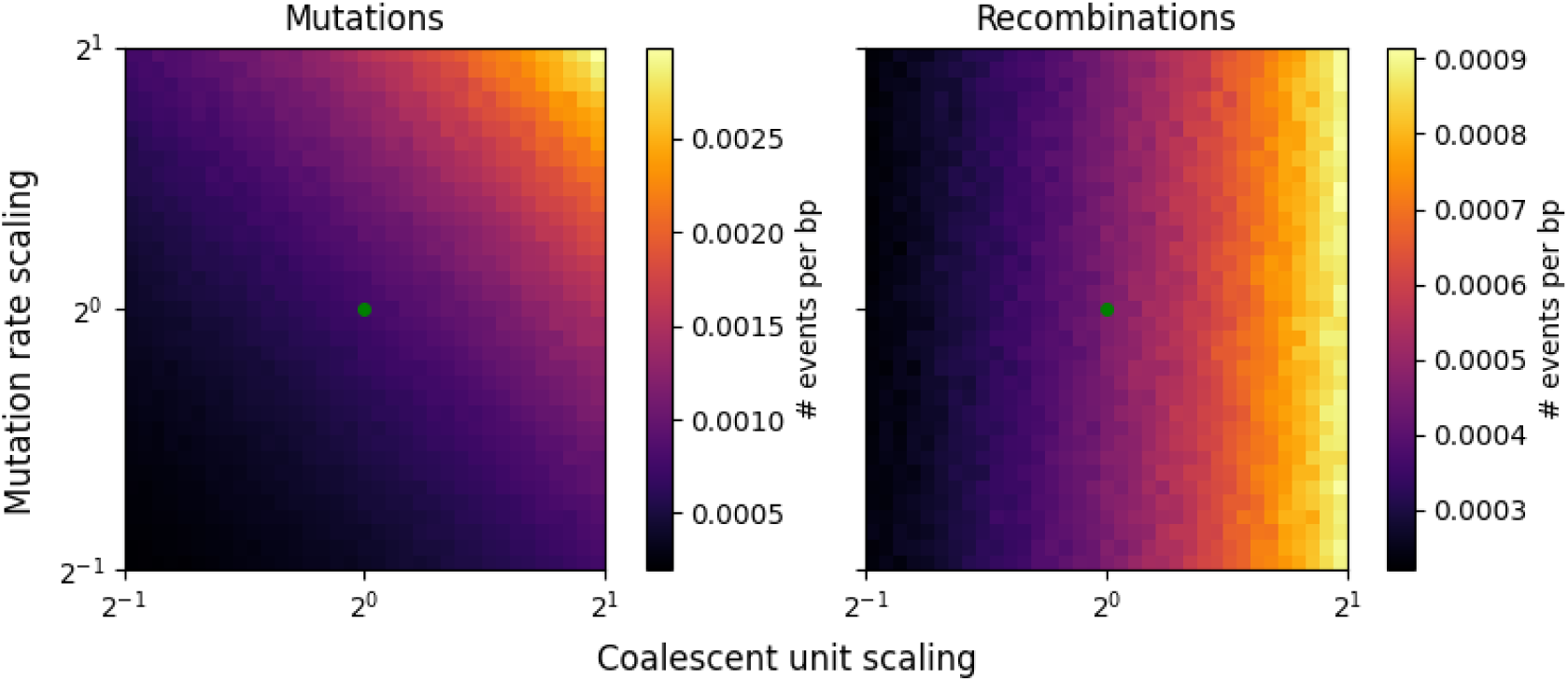
The density of mutations and recombination events across perturbations of the Zigzag stdpopsim model, for the simulations shown in Figure S2.

**Figure S4:**
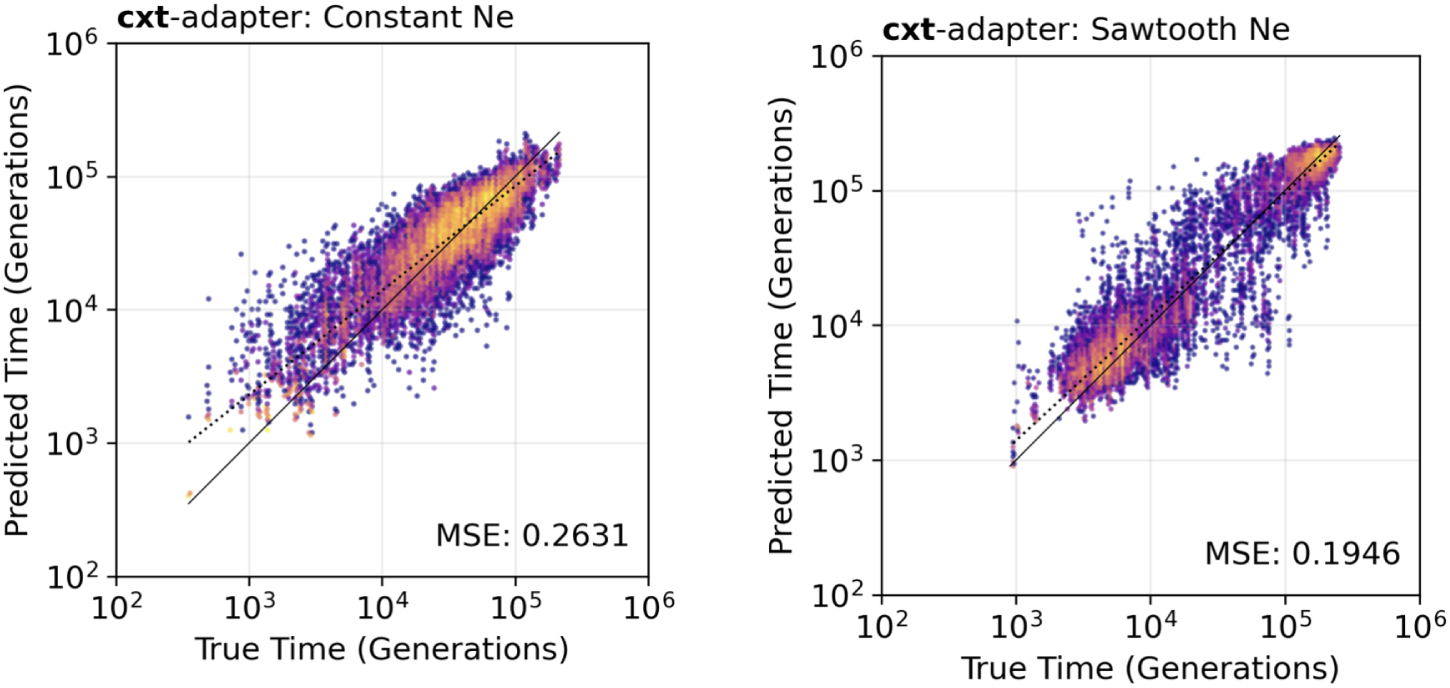
`cxt`s broad model but fine-tuned with a low-parameter adapter to account for small sample sizes. On the left, side `cxt`s is demonstrated on a constant demography, while on the right the sawtooth scenario is presented.

**Figure S5:**
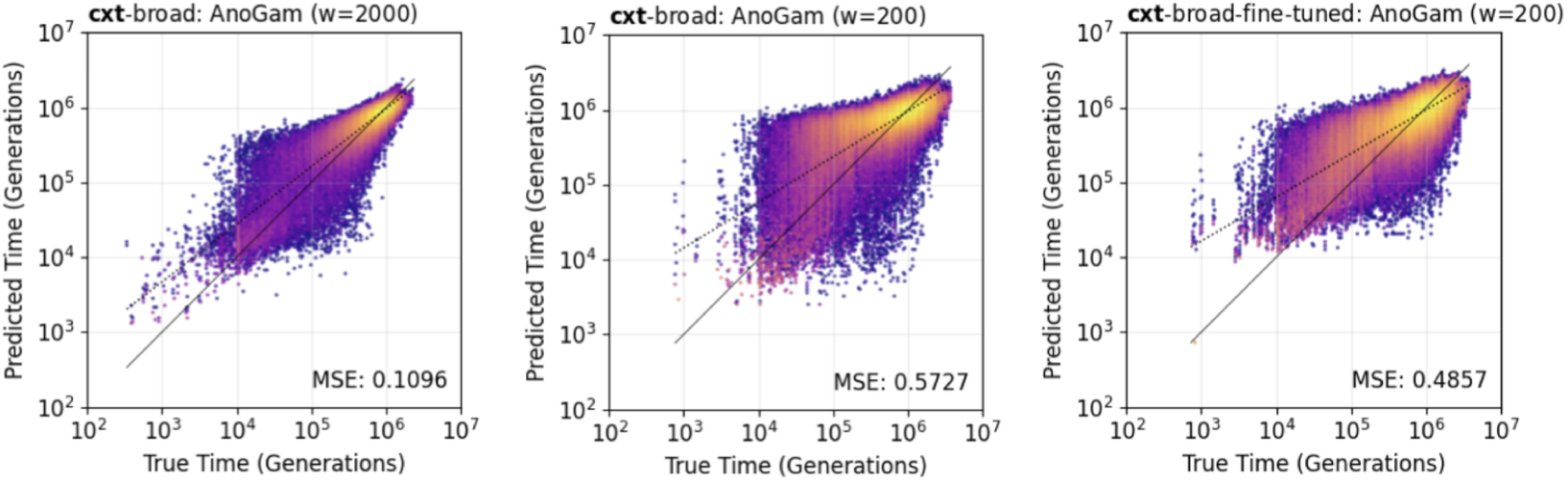
`cxt`s broad model (left) on stdpopsim AnoGam parameterization at window resolution 2 Kb, in the middle at 0.2 Kb and after fine-tuning on large stdpopsim species larger than 100 k *N_e_*.

**Figure S6:**
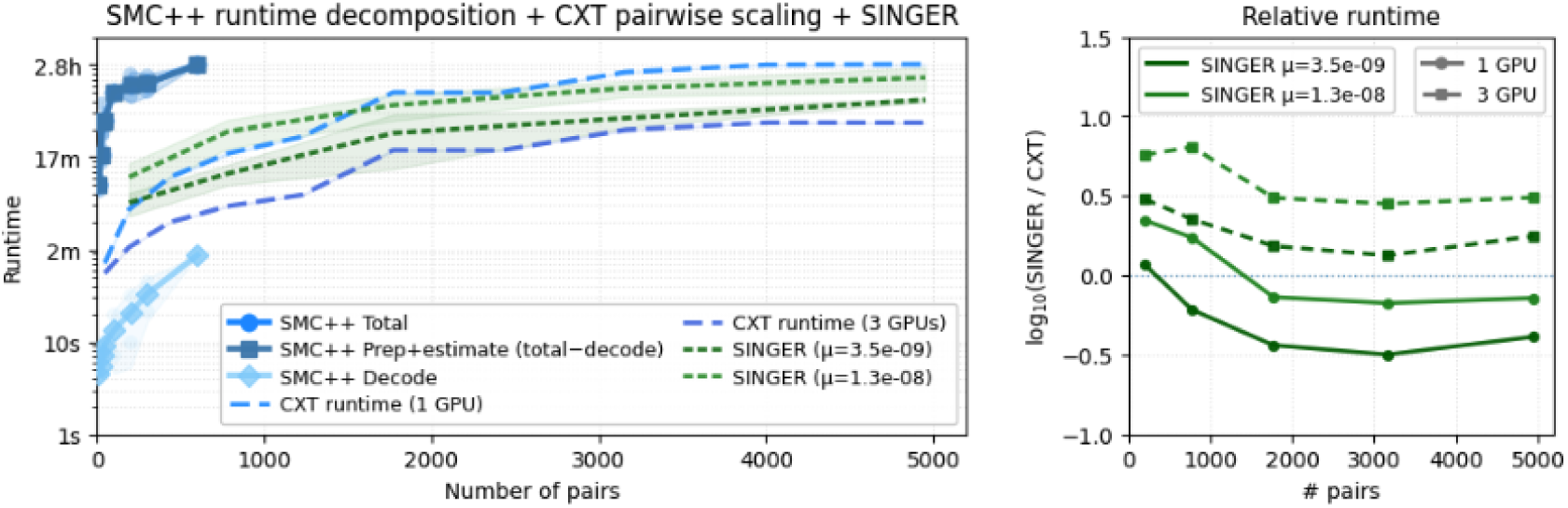
Wall-clock runtime (log scale) is shown as a function of the number of inferred haplotype pairs for SMC++ (solid; decomposed into total, preprocessing+estimate, and decoding posterior), `cxt` (dashed blue; 1 to 3 GPUs), and `Singer` (dashed green; two mutation-rate settings). Right panel shows the fold difference between `Singer` and `cxt` with respect to recombination rate or number of GPUs.

**Figure S7:**
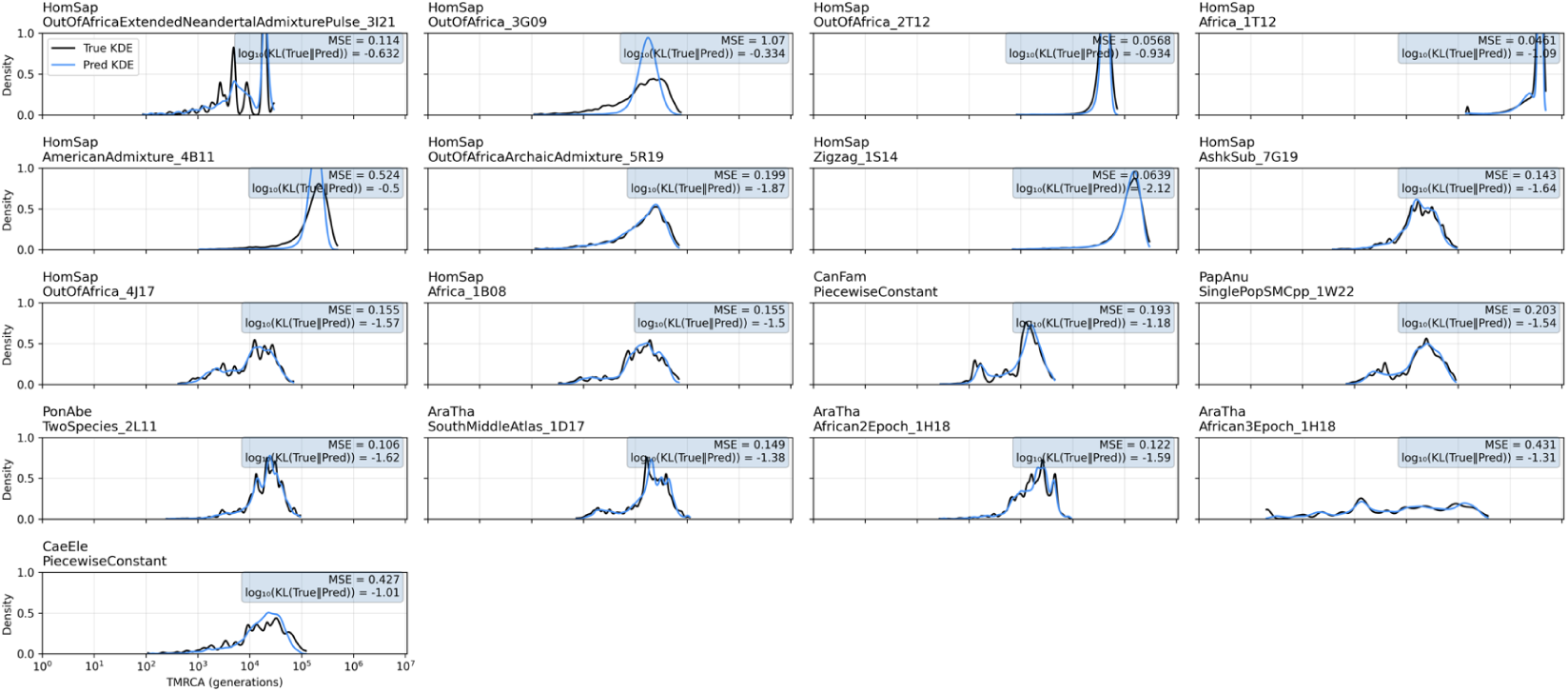
Evaluation of inferred marginal coalescence distributions (dashed line) against the true distributions (shaded line), highlighting the model’s **capacity** - its ability to distinguish scenarios based on context alone. The model generalizes to stdpopsim v0.2 simulations (**A1-F3**) with varying mutation, recombination rates, and demography. Distributions are aggregated from pairwise inferences and visualized as kernel densities. All results are from new simulations, excluded from training. Unlike the main figure, this one includes simulations with a genetic map.

**Figure S8:**
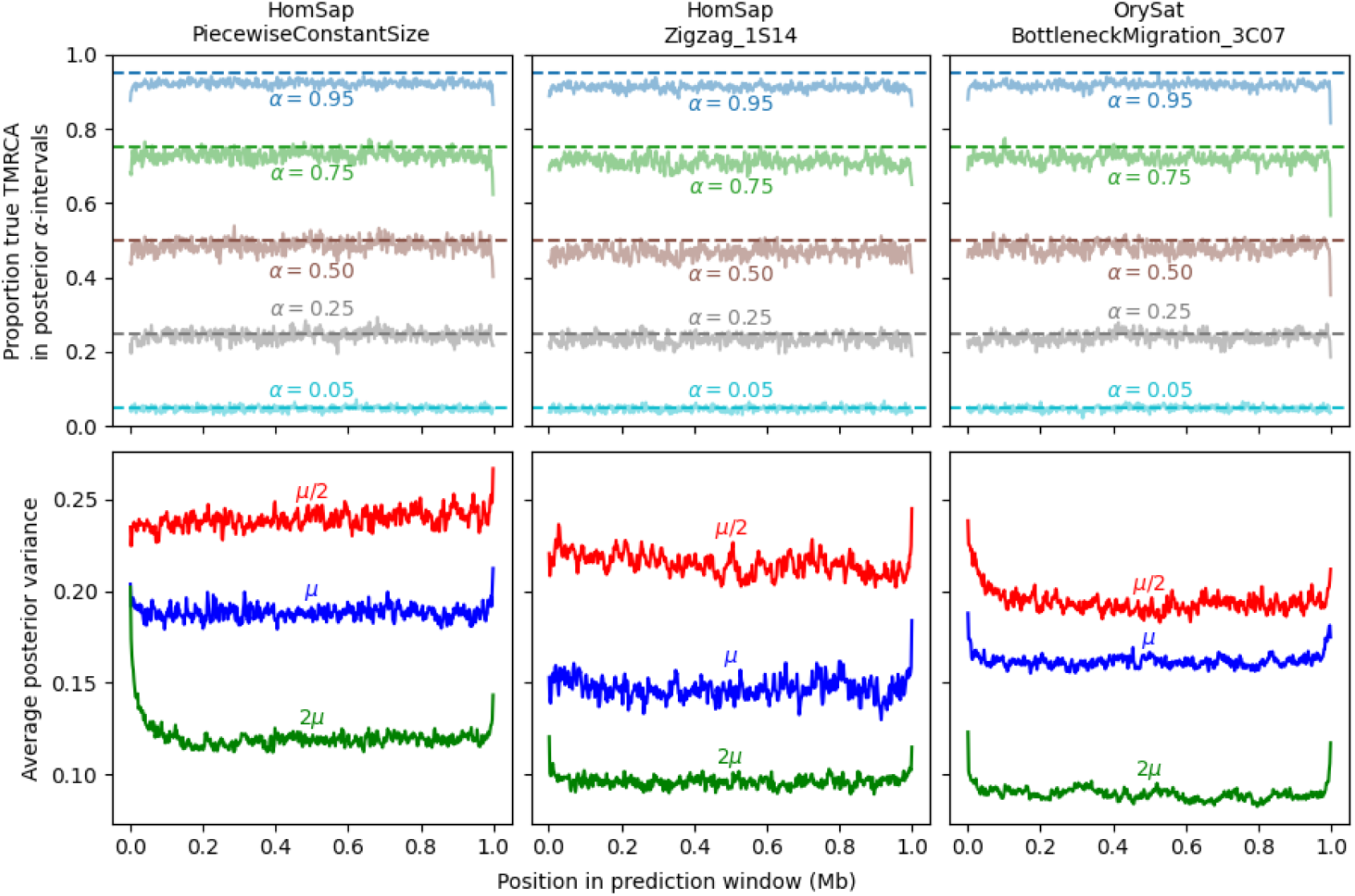
The approximate posterior distributions sampled from by `cxt` give well-calibrated estimates of uncertainty for predicted TMRCAs, across a range of scenarios. Top row: the proportion of true TMRCAs that fall within credibility intervals of a given width (e.g. a given proportion of posterior mass), as a function of position in the 1Mb prediction frame. Credibility intervals are calculated from quantiles of 100 samples from the approximate posterior for each pivot pair, and the proportion covering the true TMRCAs are calculated over 1000 independent pivot pairs (e.g. from distinct genealogical simulations). The expected proportions (for an exact posterior) are shown as dashed lines. Empirically, the approximate posteriors are asymptotically well-calibrated, although slightly over-concentrated for larger interval widths (likely due to the time discretization used by `cxt`). The decrease in coverage for the last 2Kb window is due to a tendency of the LLM to introduce large jumps in TMRCA in this window. Bottom row: the posterior variance, averaged over 1000 independent pivot pairs, as a function of position in the prediction frame and mutation rate (where `µ` stands for the species’ mutation rate in `stdpopsim`). As mutational information accrues, the posteriors become more precise; and lose precision towards the boundaries of the prediction frame where local context has been clipped.

**Figure S9:**
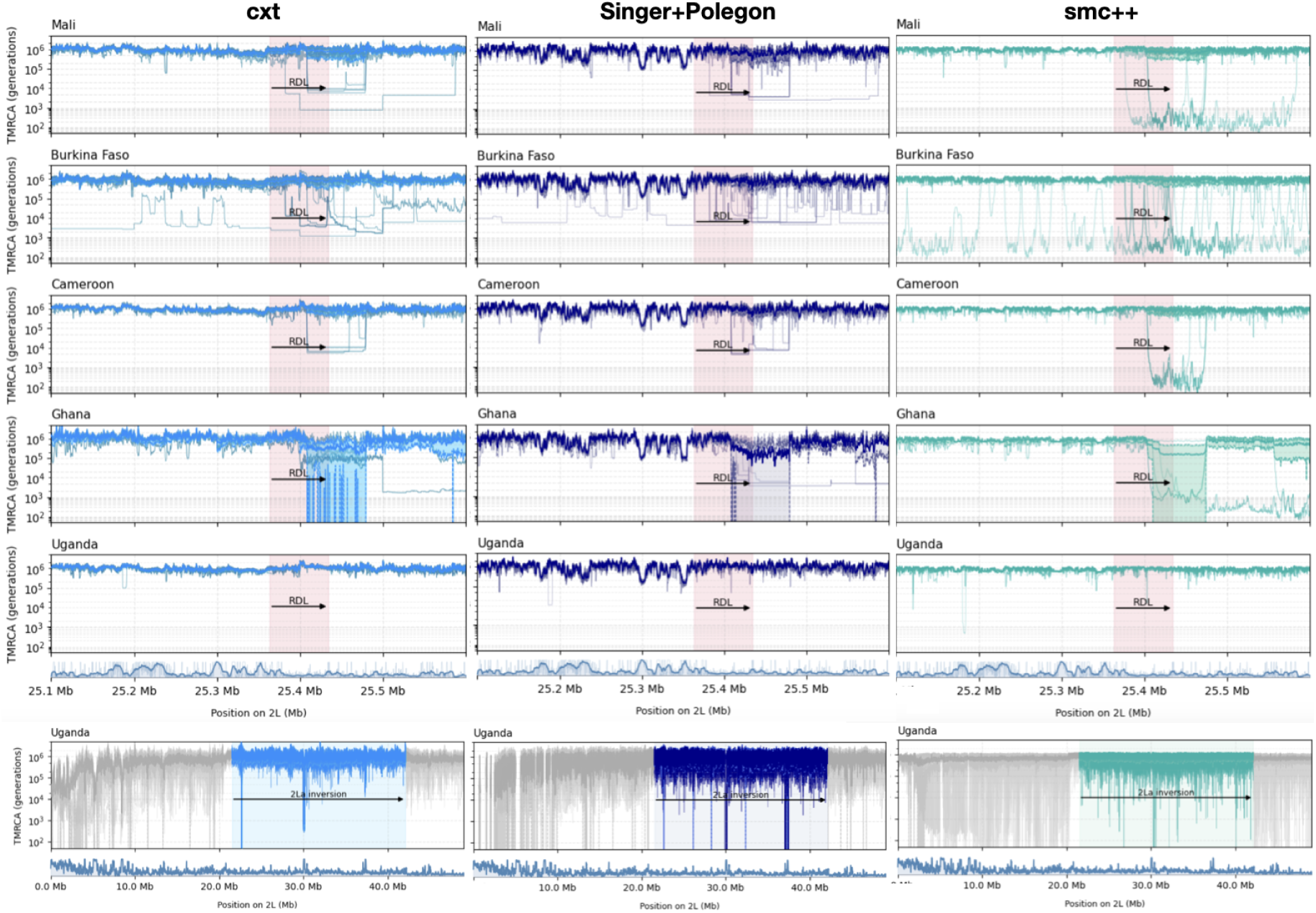
Inference of coalescent-time landscapes in the Ag1000G *A. gambiae* dataset across five African populations (Upper panel). For Burkina Faso, Mali, Cameroon, and Uganda, we analyze 25 focal pairs per population; for Ghana, we analysed only five diploid individuals. Panels show the *Rdl* region (highlighted in red), with estimates from `cxt` (left), `Singer+Polegon` (middle), and `SMC++` (right). Light-blue curves indicate per-pair inferences. The lower two panels show coalescent-time landscapes for the entirety of chr 2L for Uganda samples. Missing data density is shown beneath both the upper and lower panels as a genome browser track.

**Figure S10:**
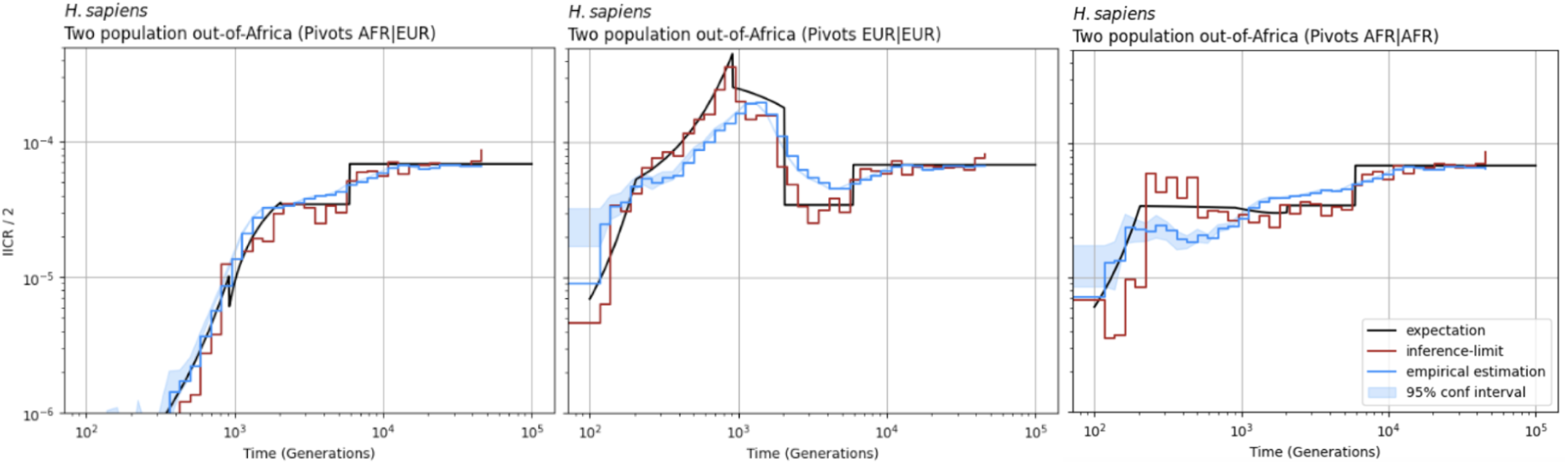
Inverse-instantaneous coalescence rate calculation of a two population out-of-Africa demogra-phy for *H. sapiens* using (**left**) only African Samples, (**middle**) only European or (**right**) a mixture of African and European samples for cross-coalescence rate estimation. The inference of pairwise-coalescence events leads to the implicit inference of demography estimates through the marginal coalescence distribu-tion assuming coalescence occurs as a Poisson process (see methods). For each scenario 10 Mb and 25 diploid samples have been used to achieve resolution throughout the specified time-windows.

**Figure S11:**
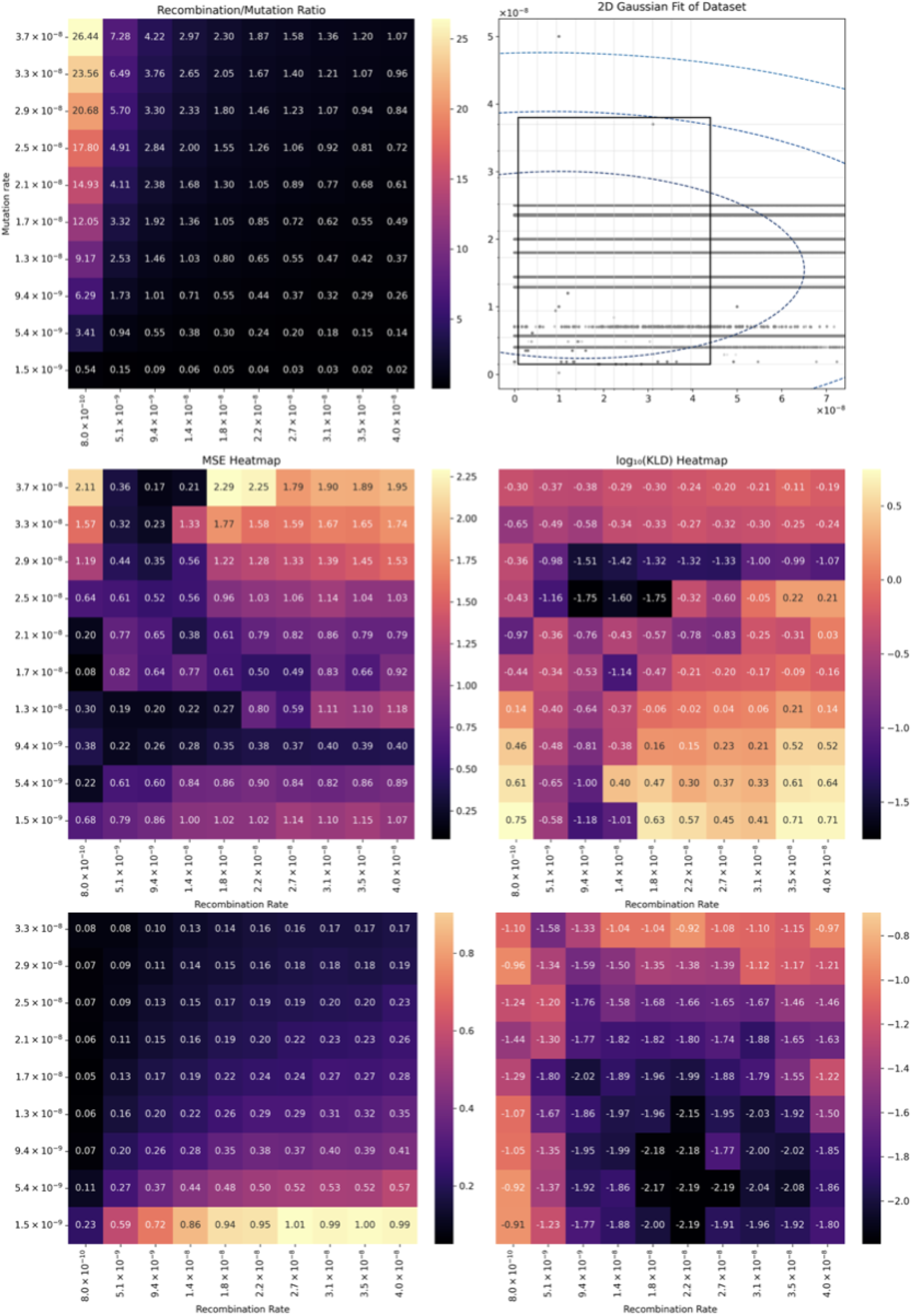
Interpretation and limitations of `cxt` by testing various recombination to mutation ratios top left. The ratios have been chosen in such way, that most of stdpopsim v0.2 with the exception of some outliers fall into this rectanlge (top right). The middle heatmaps, show MSE and KLD values of `cxt` inferences (uncalibrated) over the entire grid - while bottom heatmaps show mutation rate calibrated metrics.

**Figure S12:**
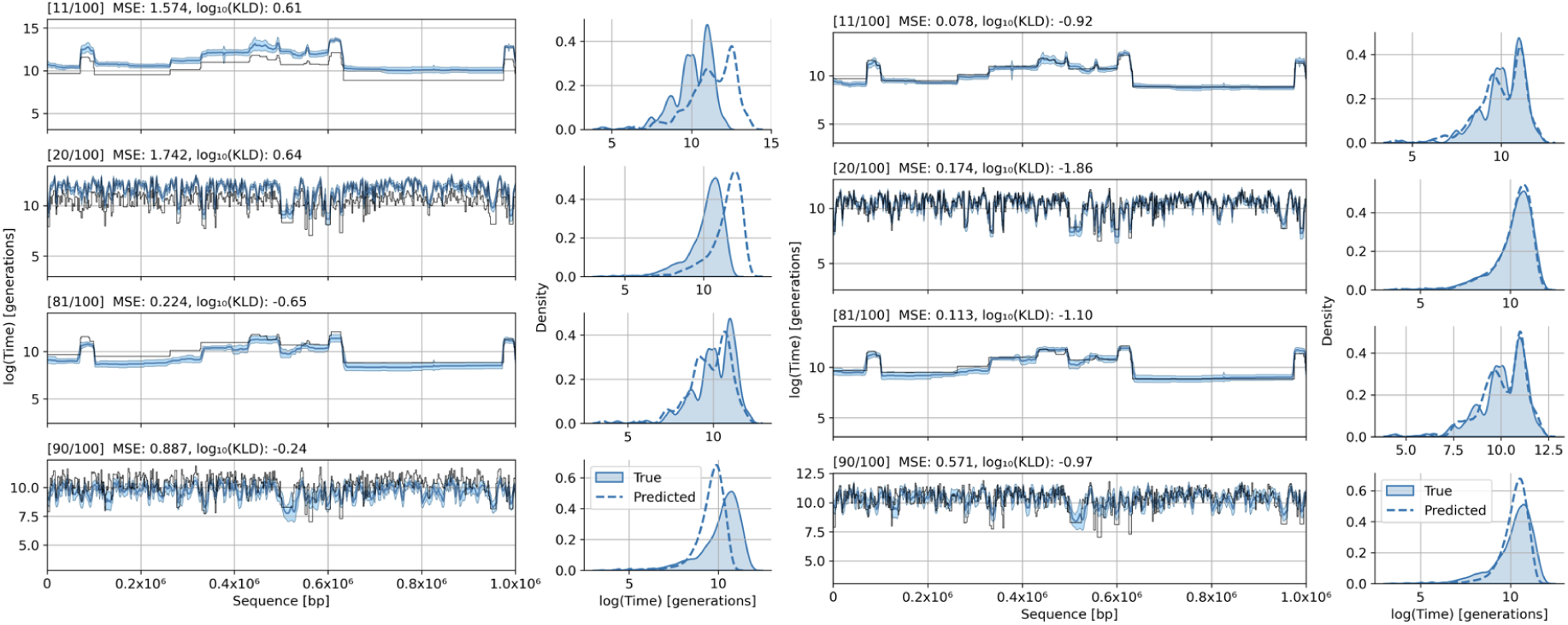
Interpretation and limitations of `cxt` zoomed into four cases of the Supplementary Figures S11. The figures are chosen to be representative for each of the corners of the heatmap of Figure S11, starting from top left to bottom right, with MSE and KLD indicated in the subtitles. These plots themselves show the TMRCA along the sequence, while both right sides show the marginal distribution, true and inferred, respectively. The left four panels are uncalibrated and the right four after calibration.

**Figure S13:**
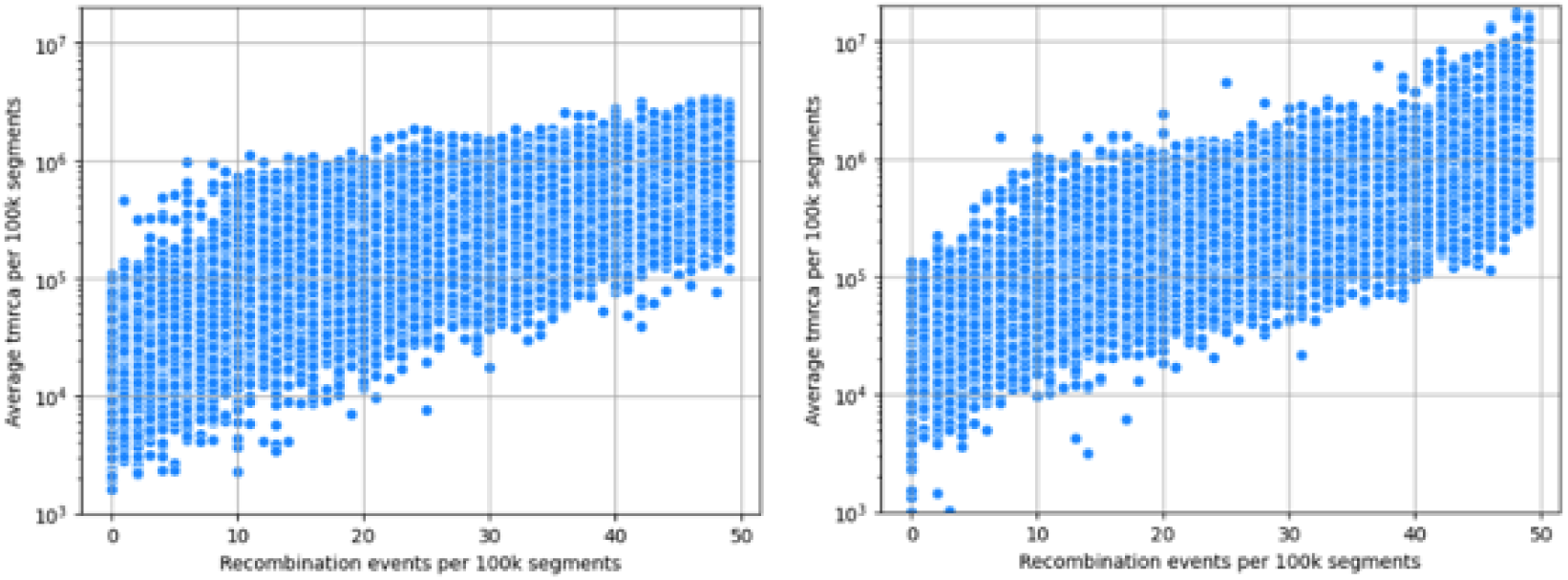
Comparison of the average TMRCA (with generation time of 28 years) for adjacent 100 Kb windows and the average number of recombination breakpoints for 25 pivot pairs using the inferences from the 1000 Genomes Project for chromosome 2 (**left**) and chromosome 6 (**right**).

## References

R. Elliott and V. Ramakrishna (1956). “Insecticide resistance in Anopheles gambiae Giles”. In: Nature 177.4507, pp. 532–533.

G. Davidson and J. Hamon (1962). “A case of dominant dieldrin resistance in Anopheles gambiae Giles”. In: Nature 196.4858, pp. 1012–1012.

J. Kingman (1982). “The coalescent”. In: Stochastic Processes and their Applications 13.3, pp. 235–248. doi: 10.1016/0304-4149(82)90011-4.

R. R. Hudson (1983). “Properties of a neutral allele model with intragenic recombination”. In: Theoretical Population Biology 23.2, pp. 183–201. doi: 10.1016/0040-5809(83)90013-8.

R. C. Griffiths (1991). “The Two-Locus Ancestral Graph”. In: Institute of Mathematical Statistics Lecture Notes - Monograph Series, pp. 100–117. doi: 10.1214/lnms/1215459289.

R. H. Ffrench-Constant et al. (1993). “A point mutation in a Drosophila GABA receptor confers insecticide resistance”. In: Nature 363.6428, pp. 449–451.

M. Thompson, J. Steichen, and R. Ffrench-Constant (1993). “Conservation of cyclodiene insecticide resistance-associated mutations in insects”. In: Insect molecular biology 2.3, pp. 149–154.

R. Griffiths and P. Marjoram (1996). “Ancestral Inference from Samples of DNA Sequences with Recombination”. In: Journal of Computational Biology 3.4, pp. 479–502. doi: 10.1089/cmb.1996.3.479.

R. C. Griffiths and P. Marjoram (1997). “An ancestral recombination graph”. In: Progress in population genetics and human evolution, pp. 257–270.

P. W. Hedrick (1998). “Balancing selection and MHC”. In: Genetica 104.3, pp. 207–214.

C. Wiuf and J. Hein (1999). “Recombination as a point process along sequences”. In: Theoretical Population Biology 55.3, pp. 248–259. doi: 10.1006/tpbi.1998.1403.

M. A. Beaumont, W. Zhang, and D. J. Balding (2002). “Approximate Bayesian Computation in Population Genetics”. In: Genetics 162.4, pp. 2025–2035. doi: 10.1093/genetics/162.4.2025.

D. M. Swallow (2003). “Genetics of lactase persistence and lactose intolerance”. In: Annual review of genetics 37.1, pp. 197–219.

T. Bersaglieri et al. (2004). “Genetic signatures of strong recent positive selection at the lactase gene”. In: The American Journal of Human Genetics 74.6, pp. 1111–1120.

W. Du et al. (2005). “Independent mutations in the Rdl locus confer dieldrin resistance to Anopheles gambiae and An. arabiensis”. In: Insect molecular biology 14.2, pp. 179–183.

G. A. McVean and N. J. Cardin (2005). “Approximating the coalescent with recombination”. In: Philo-sophical Transactions of the Royal Society B: Biological Sciences 360.1459, pp. 1387–1393. doi: 10.1098/rstb.2005.1673.

S. A. Tishkoff et al. (2007). “Convergent adaptation of human lactase persistence in Africa and Europe”. In: Nature genetics 39.1, pp. 31–40.

Y. Itan et al. (2009). “The origins of lactase persistence in Europe”. In: PLoS computational biology 5.8, e1000491.

G. P. Consortium et al. (2010). “A map of human genome variation from population scale sequencing”. In: Nature 467.7319, p. 1061.

H. Li and R. Durbin (2011). “Inference of human population history from individual whole-genome sequences”. In: Nature 475.7357, pp. 493–496. doi: 10.1038/nature10231.

B. M. Peter, E. Huerta-Sanchez, and R. Nielsen (2012). “Distinguishing between selective sweeps from standing variation and from a de novo mutation”. In: PLoS Genetics.

S. Schiffels and R. Durbin (2014). “Inferring human population size and separation history from multiple genome sequences”. In: Nature genetics 46.8, pp. 919–925. doi: 10.1038/ng.3015.

A. Auton and T. Salcedo (2015). “The 1000 genomes project”. In: Assessing rare variation in complex traits: Design and analysis of genetic studies, pp. 71–85.

L. Azevedo et al. (2015). “Trans-species polymorphism in humans and the great apes is generally maintained by balancing selection that modulates the host immune response”. In: Human genomics 9.1, p. 21.

M. C. Fontaine et al. (2015). “Extensive introgression in a malaria vector species complex revealed by phylogenomics”. In: Science 347.6217, p. 1258524.

S. Boitard et al. (2016). “Inferring Population Size History from Large Samples of Genome-Wide Molecular Data - An Approximate Bayesian Computation Approach”. In: PLOS Genetics 12.3, e1005877. doi: 10.1371/journal.pgen.1005877.

R. R. Love et al. (2016). “Chromosomal inversions and ecotypic differentiation in Anopheles gambiae: the perspective from whole-genome sequencing”. In: Molecular Ecology 25.23, pp. 5889–5906.

A. Malaspinas et al. (2016). “A genomic history of Aboriginal Australia”. In: Nature 538.7624, pp. 207–214. doi: 10.1038/nature18299.

S. Sheehan and Y. S. Song (2016). “Deep learning for population genetic inference”. In: PLoS computational biology 12.3, e1004845.

A. gambiae 1000 Genomes Consortium et al. (2017). “Genetic diversity of the African malaria vector Anopheles gambiae”. In: Nature 552.7683, p. 96.

J. Terhorst, J. A. Kamm, and Y. S. Song (2017). “Robust and scalable inference of population history from hundreds of unphased whole-genomes”. In: Nature genetics 49.2, pp. 303–309. doi: 10.1038/ng.3748.

A. Vaswani et al. (2017). Attention Is All You Need. doi: 10.48550/ARXIV.1706.03762.

A. D. Kern and D. R. Schrider (2018). “diploS/HIC: an updated approach to classifying selective sweeps”. In: G3: Genes, Genomes, Genetics 8.6, pp. 1959–1970.

P. F. Palamara et al. (2018). “High-throughput inference of pairwise coalescence times identifies signals of selection and enriched disease heritability”. In: Nature genetics 50.9, pp. 1311–1317. doi: 10.1038/s41588-018-0177-x.

A. Radford and K. Narasimhan (2018). “Improving Language Understanding by Generative Pre-Training”. In: url: https://api.semanticscholar.org/CorpusID:49313245.

L. Flagel, Y. Brandvain, and D. R. Schrider (2019). “The Unreasonable Effectiveness of Convolutional Neural Networks in Population Genetic Inference”. In: Molecular Biology and Evolution 36.2, pp. 220–238. doi: 10.1093/molbev/msy224.

F. Jay, S. Boitard, and F. Austerlitz (2019). “An ABC Method for Whole-Genome Sequence Data: Inferring Paleolithic and Neolithic Human Expansions”. In: Molecular Biology and Evolution 36.7, pp. 1565–1579. doi: 10.1093/molbev/msz038.

J. Kelleher et al. (2019). “Inferring whole-genome histories in large population datasets”. In: Nature Genetics 51.9, pp. 1330–1338. doi: 10.1038/s41588-019-0483-y.

A. Radford et al. (2019). “Language Models are Unsupervised Multitask Learners”. In.

L. Speidel et al. (2019). “A method for genome-wide genealogy estimation for thousands of samples”. In: Nature Genetics 51.9, pp. 1321–1329. doi: 10.1038/s41588-019-0484-x.

L. Torada et al. (2019). “ImaGene: a convolutional neural network to quantify natural selection from genomic data”. In: BMC Bioinformatics 20.9, p. 337. doi: 10.1186/s12859-019-2927-x.

J. R. Adrion, J. G. Galloway, and A. D. Kern (2020). “Predicting the landscape of recombination using deep learning”. In: Molecular biology and evolution 37.6, pp. 1790–1808.

C. Battey, P. L. Ralph, and A. D. Kern (2020). “Predicting geographic location from genetic variation with deep neural networks”. In: eLife 9, e54507. doi: 10.7554/eLife.54507.

C. S. Clarkson et al. (2020). “Genome variation and population structure among 1142 mosquitoes of the African malaria vector species Anopheles gambiae and Anopheles coluzzii”. In: Genome research 30.10, pp. 1533–1546.

X. Grau-Bové, et al. (2020). “Evolution of the insecticide target Rdl in African Anopheles is driven by interspecific and interkaryotypic introgression”. In: Molecular Biology and Evolution 37.10, pp. 2900–2917.

T. P. P. Sellinger et al. (2020). “Inference of past demography, dormancy and self-fertilization rates from whole genome sequence data”. In: PLOS Genetics 16.4, e1008698. doi: 10.1371/journal.pgen.1008698.

C. Battey, G. C. Coffing, and A. D. Kern (2021). “Visualizing population structure with variational autoencoders”. In: G3 11.1, jkaa036.

T. Sanchez et al. (2021). “Deep learning for population size history inference: Design, comparison and combination with approximate Bayesian computation”. In: Molecular Ecology Resources 21.8, pp. 2645–2660. doi: 10.1111/1755-0998.13224.

J. Schmidhuber (2022). Annotated History of Modern AI and Deep Learning. doi: 10.48550/arXiv.2212.11279.

K. Korfmann, O. E. Gaggiotti, and M. Fumagalli (2023). “Deep Learning in Population Genetics”. In: Genome Biology and Evolution 15.2, evad008. doi: 10.1093/gbe/evad008.

Z. Mo and A. Siepel (2023). “Domain-adaptive neural networks improve supervised machine learning based on simulated population genetic data”. In: PLoS Genetics 19.11, e1011032.

J. N. Saada et al. (2023). “Inference of Coalescence Times and Variant Ages Using Convolutional Neural Networks”. In: Molecular Biology and Evolution 40.10, msad211. doi: 10.1093/molbev/msad211.

R. Schweiger and R. Durbin (2023). Ultra-fast genome-wide inference of pairwise coalescence times. doi: 10.1101/2023.01.06.522935.

C. C. Smith and A. D. Kern (2023). “disperseNN2: a neural network for estimating dispersal distance from georeferenced polymorphism data”. In: BMC bioinformatics 24.1, p. 385.

C. C. Smith et al. (2023). “Dispersal inference from population genetic variation using a convolutional neural network”. In: Genetics 224.2, iyad068.

J. Su, et al. (Nov. 2023). RoFormer: Enhanced Transformer with Rotary Position Embedding. arXiv:2104.09864. doi: 10.48550/arXiv.2104.09864.

L. S. Whitehouse and D. R. Schrider (2023). “Timesweeper: accurately identifying selective sweeps using population genomic time series”. In: Genetics 224.3, iyad084.

B. C. Zhang et al. (2023). “Biobank-scale inference of ancestral recombination graphs enables genealogical analysis of complex traits”. In: Nature Genetics 55.5, pp. 768–776.

Y. Deng, R. Nielsen, and Y. S. Song (2024). Robust and Accurate Bayesian Inference of Genome-Wide Genealogies for Large Samples. doi: 10.1101/2024.03.16.585351.

K. Korfmann et al. (2024). “Simultaneous Inference of Past Demography and Selection from the Ancestral Recombination Graph under the Beta Coalescent”. In: Peer Community Journal 4. doi: 10.24072/pcjournal.397.

T. Sellinger, F. Johannes, and A. Tellier (2024). “Improved inference of population histories by integrating genomic and epigenomic data”. In: eLife 12. doi: 10.7554/eLife.89470.2.

J. Terhorst (2024). “Accelerated Bayesian inference of population size history from recombining sequence data”. In: bioRxiv: The Preprint Server for Biology, p. 2024.03.25.586640. doi: 10.1101/2024.03.25.586640.

G. Benegas et al. (2025). “A DNA language model based on multispecies alignment predicts the effects of genome-wide variants”. In: Nature Biotechnology, pp. 1–6.

G. Brixi et al. (2025). “Genome modeling and design across all domains of life with Evo 2”. In: bioRxiv, pp. 2025–02.

Y. Deng, Y. S. Song, and R. Nielsen (2025). “A General Framework for Branch Length Estimation in Ancestral Recombination Graphs”. In: bioRxiv, pp. 2025–02.

A. L. Fortier and J. K. Pritchard (2025). “Ancient trans-species polymorphism at the major histocompat-ibility complex in primates”. In: Elife 14, RP103547.

J. Min et al. (2025). “Neural posterior estimation for population genetics”. In: bioRxiv, pp. 2025–12.

